# Evidence for role of transketolase function in the maintenance of pyridine nucleotide levels in *Escherichia coli*

**DOI:** 10.1101/2023.03.15.532724

**Authors:** A. Vimala, R. Harinarayanan

## Abstract

The transketolase (Tkt) activity provides reversible link between glycolysis and pentose phosphate pathway (PPP). Depending on the metabolic flux, it can catalyse synthesis of glycolytic intermediates, fructose-6-phosphate and glyceraldehyde-3-phosphate from xylulose-5-P and ribose-5-P (PPP intermediates) and synthesis of xylulose-5-P from the above mentioned glycolytic intermediates. Using *E. coli*, we addressed the physiological significance of this metabolic flexibility by studying the growth phenotypes and metabolic changes associated with depletion of transketolase activity and the genetic changes or growth conditions that rescued the growth phenotypes. Tkt function was needed for cell growth when glucose was catabolized solely through Oxidative-PPP. Under gluconeogenic growth conditions, either transketolase or UdhA transhydrogenase was needed for growth. Cells depleted of Tkt activity were more sensitive than wild type to genetic changes that perturb pyridine cofactor levels. In LB medium, Tkt function was needed to prevent growth arrest from the accumulation of ribose- 5-P and possibly other pentose phosphates. In cell free extracts, the activity of Zwf and Gnd enzymes that support NADPH synthesis was inhibited by ribose-5-P. These results suggested, Tkt function played an important role in the maintenance of pyridine cofactor pool and this was confirmed by quantification. Metabolomic changes associated with transketolase depletion supported the genetic data.

## Introduction

Transketolase function is conserved across life forms, seen in bacteria, archaea and animalia. *E. coli* codes for two transketolase isoforms namely TktA and TktB (Josephson and Fraenkel 1969; Iida, et al. 1993), which catalyses reversible transfer of ketol group between multiple donor and acceptor substrates (Sprenger, 1995). The two isoforms have 75% amino acid identity. In *E. coli*, transketolase activity catalyses the following reactions, (i) transfer of ketol group between glycolytic intermediates fructose-6-P (donar) and glyceraldehyde-3-P (acceptor) to generate xylulose-5-P and erythrose-4-P and (ii) transfer of ketol group between intermediates of the pentose phosphate pathway, xylulose-5-P (donar) and ribose-5-P (acceptor) to synthesize glyceraldehyde-3-P and sedoheptolose-7-P and these can be further converted to fructose-6-P and erythrose-4-P by the action of transaldolase. Since transketolase activity is essential for synthesis of erythrose-4-P, the precursor for aromatic amino acids and pyridoxal phosphate (Vitamin B6) synthesis (Zhao and Winkler, 1994) supplementation of aromatic amino acids and pyridoxine was required for growth of ΔtktA ΔtktB strain in minimal medium. Expression of the *tktB* gene is positively regulated by RpoS and (p)ppGpp, hence the ppGpp^0^ Δ*tktA* mutant phenocopied growth properties of Δ*tktA* Δ*tktB* strain (Harinarayaan, et. al. 2008) The transketolase deficient strain was reported to be sensitive to presence of pentose sugars under gluconeogenic growth conditions, however, these studies were carried out using “leaky” transketolase mutants (Josephson and Fraenkel, 1974). We observed the Δ*tktA* Δ*tktB* strain did not grow in LB, which is a rich medium with constituents that are not entirely characterized. Furthermore, supplementation of aromatic amino acids, shikimic acid and pyridoxal phosphate failed to restore growth in LB medium. Hence, growth defect of ΔtktA ΔtktB double mutant in LB was unlikely to result from the loss of erythrose-4-P synthesis, the metabolic precursor needed for synthesis of aromatic amino acids, aromatic vitamins and vitamin B6. To understand the physiological basis for growth defect in LB medium, we identified genetic changes and growth conditions that rescued the growth defect.

Genetic characterization of the mutants together with biochemical measurements indicated the growth defect of Δ*tktA* Δ*tktB* mutant in LB medium and its sensitivity to pentose sugars in gluconeogenic medium (Josephson and Fraenkel, 1974) are related. Our results indicate, the two phenotypes follow from the perturbation of NADPH/NADH metabolism. Genetic data indicated the sensitivity to ribose (which we have used as a representative pentose sugar) seen during gluconeogenic growth conditions resulted from reduction in NADH pool. The growth defect in LB was associated with depletion of NAD^+^, NADH and NADPH, although the growth defect followed primarily from NADPH depletion. Based on our results, we propose that, by modulating the metabolic flux between glycolysis and pentose phosphate pathway, the transketolase function contributes to the maintenance of cellular NADH/NADPH pool and redox homeostasis.

## Results

### Use of conditional replication plasmid for maintenance of transketolase deficient strain

Multiple attempts were made to introduce Δ*tktB*::kan allele into the MG1655 Δ*tktA*::FRT strain by phage P1 transduction. However, Kan^r^ transductants could not be recovered in LB medium and LB medium supplemented with tyrosine, tryptophan and phenylalanine (or shikimic acid) and pyridoxine (data not shown). This suggested, the Δ*tktA*::FRT Δ*tktB*::kan strain may be in- viable in LB medium and transketolase function could be required for growth in LB. To test this possibility, a plasmid based conditional expression strategy was used to study the growth of Δ*tktA*::FRT Δ*tktB*::FRT strain in LB medium. The *tktB* gene was cloned into pAM34, an Amp^r^ plasmid which exhibits IPTG-dependent replication (Gil and Bouche, 1991), and the resulting construct pAM-tktB (Vimala and Harinarayanan, 2016), was introduced into the wild type strain MG1655. The chromosomal *tktA* and *tktB* genes were then inactivated in the MG1655/pAM-tktB strain by sequential transduction and removal of antibiotic marker using FLP recombinase (see methods for details). The resulting strain, AV104 was maintained in the presence of ampicillin and IPTG to support replication of plasmid pAM-tktB. If transketolase function was required for growth in LB, it was expected that the final strain, Δ*tktA*::FRT Δ*tktB*::FRT / pAM-tktB (henceforth referred as Δ*tkt/*pAM-tktB) would exhibit IPTG dependent growth in LB medium. Consistent with that, the Δ*tkt/*pAM-tktB strain exhibited growth defect only upon removal of ampicillin and IPTG in broth (Fig. 1A, i) and as well as on LB agar plate (Fig. 1A, ii). The growth kinetics in broth indicated that several round of cell division can follow after the removal of IPTG before growth arrest followed from the depletion of transketolase activity. After testing different amounts of inoculum, it was found that 1:1500 dilution of an overnight culture (grown in the presence of ampicillin and IPTG) supported growth up to A_600_ of 0.3 - 0.4 in LB (Fig. 1A, i). Throughout this study, this growth condition was used to deplete transketolase activity and carry out genetic or biochemical experiments to address the role of transketolase function under different growth conditions.

**Fig. 1.**
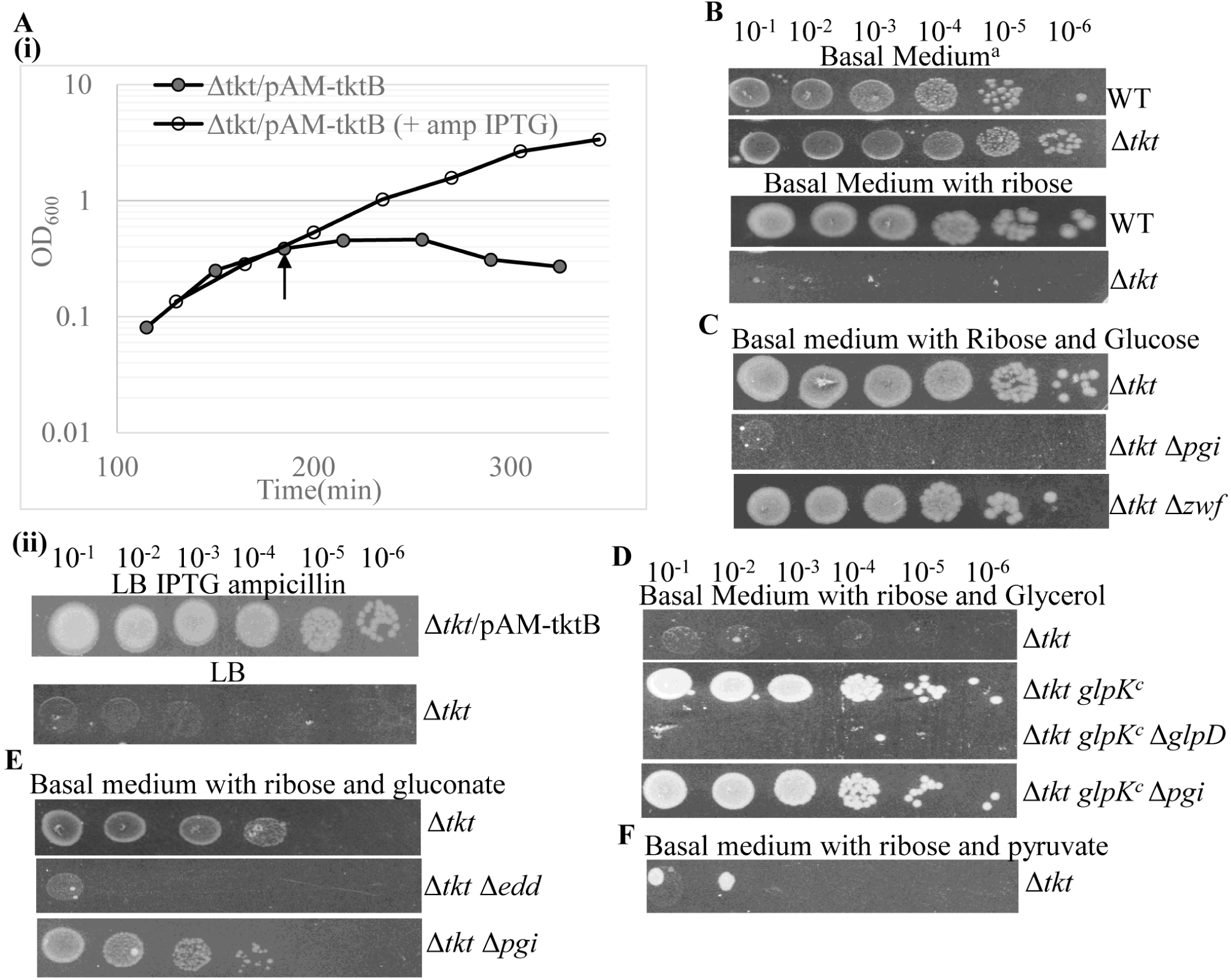
Depletion of transketolase activity conferred growth arrest in LB medium and ribose sensitivity in basal medium - effect of sugars and genetic changes on ribose sensitivity. Overnight culture of Δ*tkt* /pAM-tktB strain in LB amp IPTG (1 mM) was washed, diluted 1:1500 and sub-cultured into LB amp IPTG (1 mM) and LB and A_600_ was monitored over time [A(i)]. Cells were collected from LB culture at the time point indicated by arrow, serially diluted and spotted on LB or LB amp IPTG plates [A(ii)]. Serial dilutions of strains whose relevant genotypes are indicated were depleted for transketolase activity and spotted on basal medium and basal medium supplemented with sugars as indicated (B - F). ^a^ - minimal salts medium containing casaminoacids, tryptophan, pyridoxine and shikimic acid (see material and methods for details). Strains, WT (MG1655); Δ*tkt/*pAM-tktB (AV104); Δ*tkt* Δ*pgi/*pAM- tktB (AV108); Δ*tkt* Δ*zwf/*pAM*-*tktB (AV107); Δ*tkt glpK^c^*/pAM*-*tktB (AV148); Δ*tkt glpK^c^* Δ*glpD/*pAM*-*tktB (AV150); Δ*tkt glpK^c^* Δ*pgi/*pAM*-*tktB (AV234) and Δ*tkt* Δ*edd/*pAM*-*tktB (AV109).

### Glycolytic catabolism of glucose, gluconate and glycerol rescued the ribose sensitivity of Δ*tkt* mutant

The Δ*tktA* Δ*tktB* strain, cannot synthesize erythrose-4-P, a metabolic intermediate necessary for synthesis of aromatic amino acids, aromatic vitamins and pyridoxine (vitamin B6) (Zhao and Winkler, 1994). An early study, that used “leaky” transketolase mutants reported growth sensitivity to pentose sugars under gluconeogenic growth conditions and rescue by glucose supplementation (Josephson and Fraenkel, 1974). To find out if depletion of transketolase activity gave rise to similar phenotypes, we studied the growth response of Δ*tkt*/pAM-tktB strain and isogenic mutant derivatives of this strain to sugars following the depletion of transketolase activity. Minimal A medium (Miller,1992) containing casamino acids and supplemented with shikimic acid and pyridoxine to satisfy aromatic amino acid (primarily tryptophan which is known to be deficient in casamino acids), aromatic vitamins and vitamin B6 requirements of Δ*tkt* strain was used (henceforth referred as basal medium). Throughout this study, we have used ribose as a representative pentose sugar to induce growth arrest in transketolase depleted strain. In most instances, the phenotypes elicited by ribose was also observed following supplementation of arabinose or xylose (data not shown).

The transketolase depleted strain, but not wild type, exhibited ribose sensitivity in basal medium (Fig. 1B) and this was rescued by glucose supplementation (Fig. 1C). Since glucose can be catabolized through glycolysis and the pentose phosphate pathway (PPP), we tested the role of each pathway in growth rescue. Glucose mediated rescue of ribose sensitivity was completely lost following the inactivation of *pgi* (encoding phophoglucoisomerase that catalyse conversion of glucose-6-P to fructose-6-P) and was unaffected after inactivation of z*wf* (encoding glucose-6-P dehydrogenase, required for the Entner Doudoroff (ED)/PPP mediated catabolism of glucose). This suggested, glycolytic catabolism of glucose was necessary and sufficient for growth rescue (Fig. 1C).

We had previously reported a gain of function mutation, *glpK*^C^, from transposon insertion at the end of the *glpF* gene and which resulted in constitutive *glpK* expression (Vimala and Harinarayanan, 2016). The ribose sensitivity of transketolase depleted strain was rescued by the *glpK*^C^ allele in the presence of glycerol, while *glpK*^C^ or glycerol separately did not rescue, and the growth rescue was dependent on *glpD* gene function (Fig. 1D). These results suggested, the catabolism of glycerol to glycolytic intermediate dihydroxyacetone phosphate (DHAP) by GlpD was necessary for the growth rescue.

Gluconate supplementation partially rescued the ribose sensitivity of transketolase depleted strain in basal medium and this was, dependent on the *edd* function (Fig. 1E), indicating that catabolism of gluconate via the ED pathway to glycolytic intermediates, glyceraldehyde-3-P (Gld-3-P) and pyruvate was necessary for the rescue of ribose sensitivity. Pyruvate supplementation did rescue ribose sensitivity (Fig. 1F) and suggested sugars that feed in to the end of the glycolytic pathway cannot rescue. To test if gluconeogenic catabolism of glycerol or gluconate played a role, we introduced the Δ*pgi* mutation. The rescue of ribose sensitivity by glycerol in the presence of the *glpK*^C^ allele or gluconate was unaffected (Figs. 1D and 1E). Since ribose sensitivity of transketolase depleted strain was (i) rescued by glucose, gluconate and glycerol and dependent on *pgi*, *edd* and *glpD* functions respectively (ii) rescue by gluconate and glycerol was unaffected by the loss of *pgi* function and (iii) pyruvate could not rescue the growth defect, we propose the following. Glycolytic catabolism of the sugars was required for growth rescue, and reaction(s) shared between the catabolic pathways of glucose, gluconate and glycerol, but not pyruvate, were required for rescue. Therefore, we hypothesize that the glycolytic reaction(s) involved in conversion of glyceraldehyde-3-P to pyruvate was required for rescue of ribose sensitivity of transketolase depleted strain.

### Transketolase function was necessary for growth when glucose was catabolized solely through Oxi-PPP pathway

Unlike Δ*tkt* mutant, the Δ*pgi* Δ*tkt* double mutant, exhibited growth defect in basal medium and basal medium supplemented with glucose (Fig.2A). The Δ*pgi* single mutant grew in the basal medium with or without glucose (data not shown). The grow defect of Δ*pgi* Δ*tkt* mutant in basal medium may be attributed to the inability to synthesize ribose-5-P an essential metabolite via gluconeogenesis. Growth defect in the presence of glucose was unexpected and suggested transketolase activity was essential for growth on glucose and amino acids when the Pgi mediated catabolism of glucose was blocked, leaving glucose to be catabolized through the Oxi-PPP or ED pathways. Glucose catabolism through ED pathway is insignificant in wild type strain (Fraenkel & Levisone, 1967; Fraenkel DG, 1986; Eisenberg and Dobrogosz, 1967), but may become the primary route of glucose catabolism in the Δ*pgi* Δ*tkt* mutant. However, the Δ*tkt* Δ*pgi* Δ*edd* strain continued to exhibit growth defect in the basal glucose medium (data not shown), indicating glucose catabolism through ED pathway was not the reason for growth inhibition, and suggested to a possible role for Oxi-PPP mediated catabolism of glucose.

**Fig. 2.**
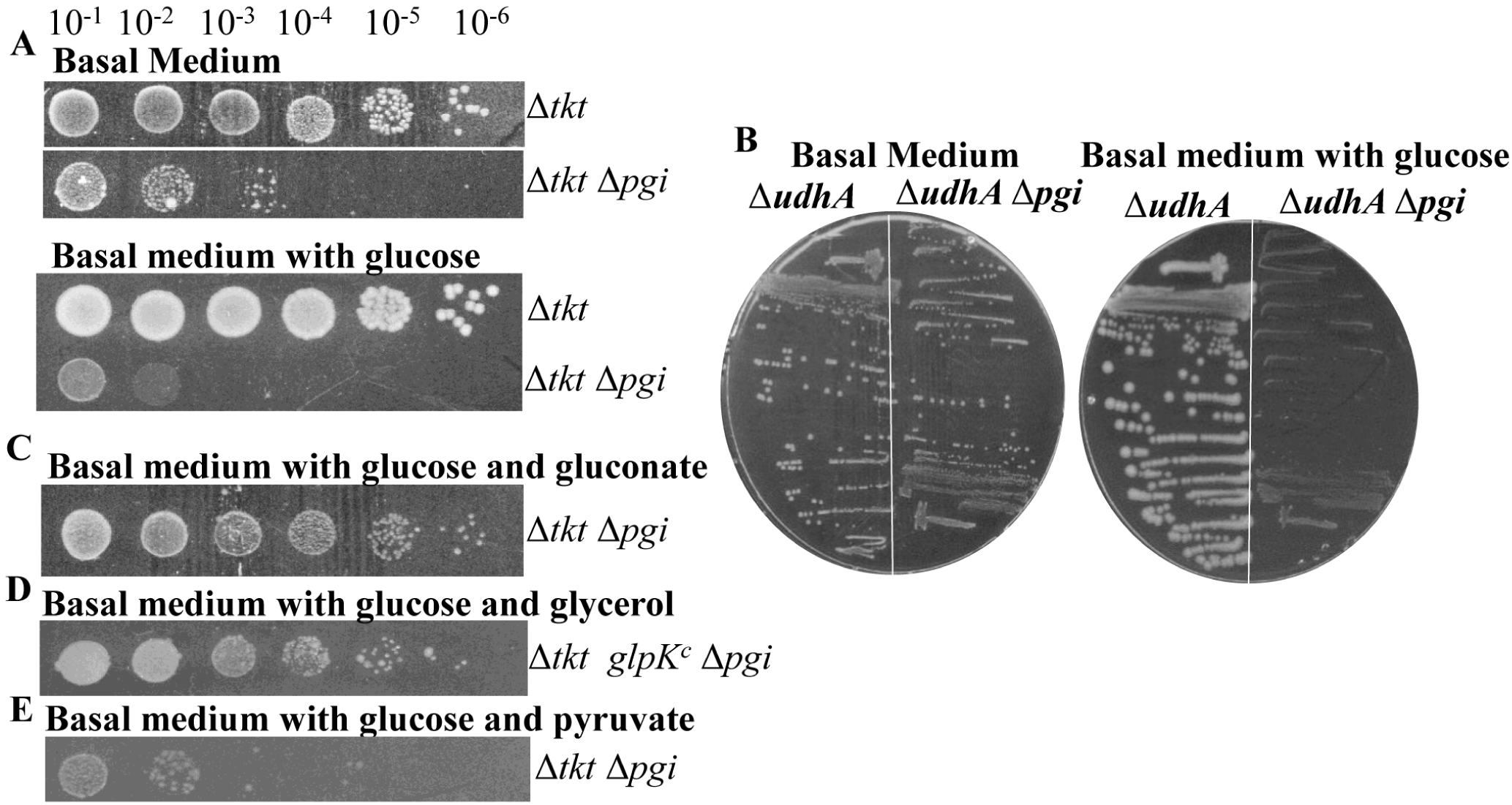
Loss of Tkt or UdhA function confers glucose sensitivity in the Δpgi background and this was rescued by glycerol or gluconate. Serial dilutions of strains whose relevant genotypes are indicated were depleted for transketolase activity and spotted (A, C - E) or streaked (B) on basal medium and in the presence of sugars as indicated. Δ*tkt/*pAM*-*tktB (AV104); Δtkt Δ*pgi/*pAM-tktB (AV108); Δ*tkt glpK^c^* Δ*pgi/*pAM*-*tktB (AV234); Δ*udhA* (AV198) and Δ*udhA* Δ*pgi* (AV309).

To decipher the basis for Δ*tkt* Δ*pgi* growth defect, we considered phenotypes reported for the Δ*pgi* mutant in glucose. Exclusive catabolism of glucose through Oxi-PPP in the Δ*pgi* mutant reduced growth rate, and this was rescued by the over-expression of the cytoplasmic transhydrogenase UdhA (Canonaco, F *et al*. 2001). Further, the growth defect of Δ*pgi* mutant was accentuated by Δ*udhA* mutation (Sauer, U *et al*. 2004, Hansen and Schönheit, 2005). Since primary role of UdhA during growth on glucose was NADH synthesis concomitant with oxidation of NADPH (Sauer, U *et al*., 2004), it was suggested, the increase in glucose catabolism by Oxi-PPP (due to the Δ*pgi* mutation) and reduced synthesis of NADH (due to the Δ*udhA* mutation) caused an increase in NADPH to NADH ratio (Canonaco, F et al. 2001). This can also explain the growth defect of Δ*pgi ΔudhA* strain in basal medium with glucose (Fig. 2B). As described below, an increase in NADPH / NADH ratio may also be expected in the Δ*pgi* Δ*tkt* mutant during growth on glucose.

In the Δ*pgi* mutant, glucose catabolism through Oxi-PPP would result in the synthesis of pentose phosphates. Since further catabolism of pentose phosphates to glycolytic intermediates, fructose-6-P and glyceraldehyde-3-P requires transketolase activity, this would be impaired in the Δ*pgi* Δ*tkt* strain. As glycolysis and Oxi-PPP are primarily sources for NADH and NADPH respectively (Sprenger, 1995; Centeno-Leija et al. 2013), increased catabolism of glucose through Oxi-PPP (due to the Δ*pgi* mutation) and inability to synthesize glycolytic intermediates from the pentose phosphates (due to the Δ*tkt* mutation) could elevate NADPH to NADH ratio. We therefore propose, when glucose was catabolized solely through Oxi-PPP, resulting in increased synthesis of NADPH, both UdhA and Tkt activities are required to maintain the NADH synthesis and prevent growth inhibition from an increase in NADPH / NADH ratio.

We tested if growth conditions or genetic changes that increase glycolytic flux can lower the NADPH/NADH ratio and rescue the growth defect of Δ*pgi* Δ*tkt* strain. Growth defect was rescued by gluconate supplementation (Fig. 2C), glycerol supplementation in the presence of *glpK*^C^ allele (Fig. 2D) but not pyruvate supplementation (Fig. 2E). Catabolism of glycerol or gluconate would increase the glyceraldehyde-3-P pool and GAPDH dependent NADH synthesis. As this is the only glycolytic step upstream of pyruvate that can support NADH synthesis, we propose it may be responsible for the growth rescue of Δ*pgi* Δ*tkt* strain by alleviating the reduction in NADPH/NADH ratio. The model we propose is, whenever there is glucose catabolism and associated NADPH synthesis through Oxi-PPP, GAPDH dependent NADH synthesis is necessary for the maintenance of NADPH/NADH balance needed for growth.

### Either transketolase or UdhA transhydrogenase activity was required for growth with amino acids as carbon source

Amino acids constiture the primary carbon source in the basal medium used here, therefore metabolism was expected to be predominantly gluconeogenic. We found that, under this growth condition, the UdhA transhydrogenase was required for growth of Δ*tkt* mutant, but not the wild type strain (Fig. 3A). Since UdhA supports NADH synthesis, it was possible the Δ*tkt* Δ*udhA* growth defect was due to a severe perturbation of NADH metabolism. Supplementation of glucose or gluconate rescued the growth defect of Δ*tkt* Δ*udhA* mutant in basal medium (Fig. 3B), and suggested, switching from gluconeogenic to glycolytic mode of metabolism rescued the growth defect of Δ*tkt* Δ*udhA* mutant. However, glycerol or glycerol in the presence of *glpK*^C^ allele failed to rescue growth defect (Fig. 3C). Interestingly, growth rescue was seen in the presence ribose (Fig. 3C). Pyruvate did not rescue growth defect either by itself or together with ribose (data not shown).

**Fig. 3.**
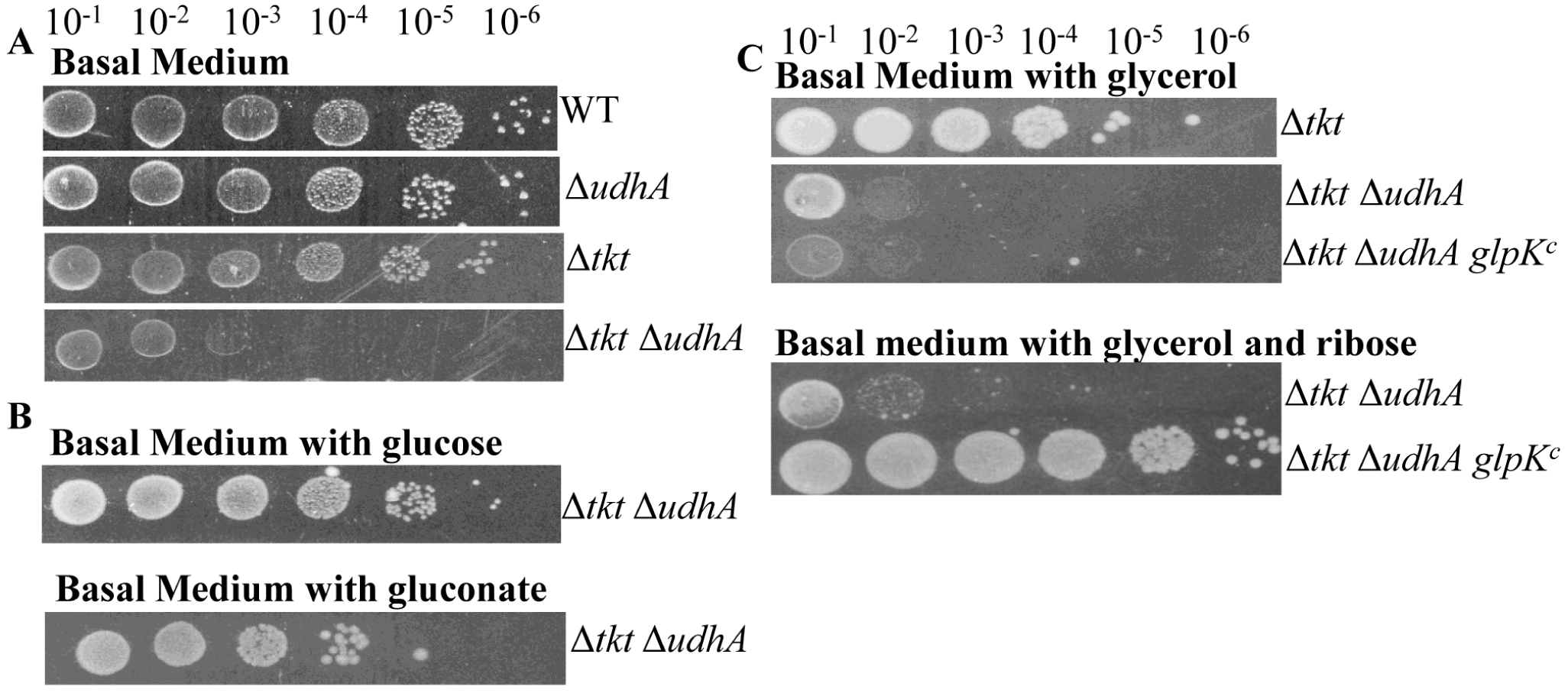
Transketolase or UdhA activity was required for growth in medium with amino acids as primary carbon source. Strains whose relevant genotypes are indicated were depleted for transketolase activity, serially diluted and spotted on the growth medium indicated (A – C). WT (MG1655), ΔudhA (AV198), Δ*tkt/*pAM*-*tktB (AV104); Δ*tkt* Δ*udhA/*pAM-tktB (AV208) and Δ*tkt* Δ*udhA glpK^c^/* pAM*-*tktB (AV187)

As the catabolic pathways of sugars that confer growth rescue (glucose, gluconate or glycerol) converge at glyceraldehyde-3-P and pyruvate supplementation did not rescue, an increase in the glycolytic flux from Gld-3-P could be important for growth rescue. The reason growth rescue by glycerol was ribose dependent could be related to inhibition of Zwf and Gnd enzyme activities by ribose-5-P (described below) and consequent increase in glycerol catabolism through glycolysis (relative to gluconeogenesis).

The model we propose is, UdhA and transketolase functions, contribute directly and indirectly to NADH synthesis during growth in basal medium. Therefore, growth defect of Δ*tkt* Δ*udhA* strain in basal medium arises from NADH depletion. When using amino acids as carbon source, (i) transketolase dependent synthesis of fructose-6-P and glyceraldehyde-3-P from the pentose phosphates support GAPDH dependent NADH synthesis (ii) UdhA supports NADH synthesis by oxidizing NADPH synthesized by Zwf and Gnd activities and (iii) at least one of the two activity was required to maintain the NADPH/NADH ratio necessary for gluconeogenic growth.

### Over-expression of PntAB transhydrogenase inhibited growth of Δtkt mutant in glucose and this was rescued by ribose

The similar growth phenotypes of Δ*pgi* Δ*udhA* and Δ*pgi* Δ*tkt* strains in the presence of glucose (Fig. 2A and 2B) and the growth defect conferred by introduction of ΔudhA mutation into Δtkt strain in basal medium (Fig. 3A), suggested that transketolase activity could be needed for maintaining NADH synthesis and the NADPH/NADH ratio under these growth conditions. We asked, if the transketolase deficient strain was differentially sensitive, as compared to wild type strain, to any other genetic perturbation known to alter NADPH/NADH metabolism. PntAB function is reported to contribute 35-45% of cellular NADPH during growth on glucose (Sauer *et al*., 2004). We found, expression of *pntAB* from plasmid severely inhibited growth of Δ*tkt* mutant, but not the wild type strain, in basal medium with glucose (Fig. 4A). On the other hand, over-expression of pntAB did not affect growth of wild type or Δ*tkt* mutant in basal medium (gluconeogenic growth) (Fig. 4B). This suggested, transketolase function could be specifically required to cope with changes in the NADPH/NADH pool during glucose catabolism. During growth on glucose, loss of transketolase function can lower the NADH pool by decreasing the synthesis of glycolytic intermediates used for NADH synthesis. The over-expression of PntAB can further lower the NADH pool by catalysing synthesis of NADPH at the expense of NADH and confer growth inhibition.

**Fig. 4.**
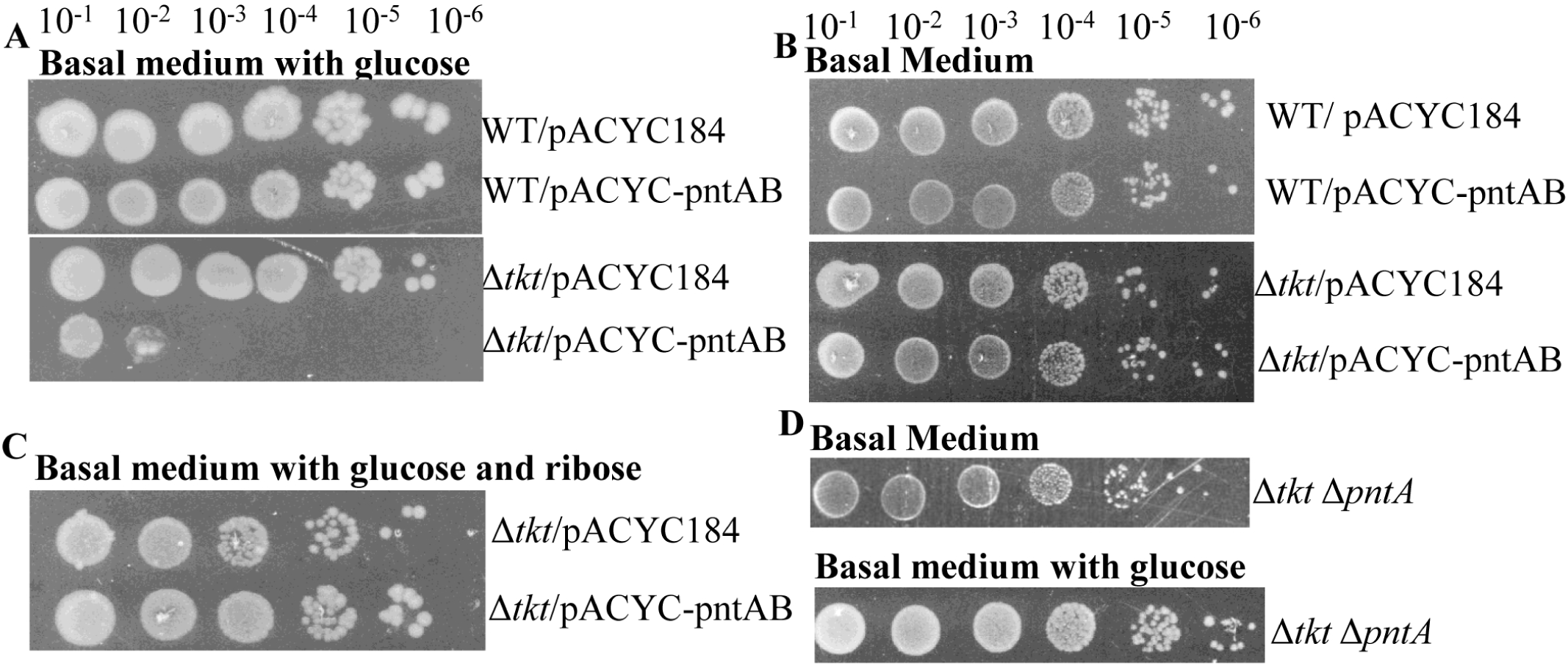
Glucose dependent growth inhibition of Δ*tkt* mutant by the over-expression of pntAB and its rescue by ribose. Strains whose relevant genotypes are indicated were depleted for transketolase activity, serially diluted and spotted in the growth media as indicated (A – D). Strains, WT/pACYC184 (AV43); WT/ pACYC-pntAB (AV45); Δ*tkt/*pAM*-tktB*/pACYC184 (AV110); Δ*tkt/*pAM*-*tktB/ pACYC-pntAB (AV112); Δ*tkt* Δ*pntA/*pAM-tktB (AV130).

Interestingly, the growth defect arising from over-expression of *pntAB* was rescued by ribose supplementation (Fig. 4C). Since catabolism of pentose phosphates via non-Oxi-PPP was blocked in the Δ*tkt* mutant, ribose supplementation could increase the intracellular concentration of ribose-5-P and possibly other pentose phosphates as well in the strain relative to wild type (this is supported by the metabolome data described below). As described in the next section, the activity of Zwf and Gnd (enzymes that support NADPH synthesis) was inhibited *in vitro* by ribose-5-P. Based on this observation, we suggest, growth inhibition due to *pntAB* over-expression in glucose followed from an increase in the cellular NADPH/NADH ratio, and the accumulation of ribose-5-P following ribose supplementation in Δ*tkt* mutant alleviated this by inhibiting Zwf and Gnd mediated NADPH synthesis and simultaneously redirecting metabolic flux into glycolysis to increase NADH synthesis.

Introducing Δ*pntA* mutation did not inhibit growth of Δ*tktA* mutant in basal medium with or without glucose (Fig. 4D). Since PntAB activity contributes to increasing the NADPH/NADH ratio, a decrease in this ratio may be expected from the loss of PntA function. This suggested, the transketolase depleted strain, in contrast to its sensitivity to increase in NADPH/NADH ratio may be tolerant to decrease in the ratio.

### Inhibition of Zwf and Gnd enzyme activity by ribose-5-P in cell free extracts

Since catabolism of pentose phosphates into glycolytic intermediates require transketolase activity, ribose supplementation can lead to the accumulation of pentose phosphates in the Δ*tkt* mutant. This is supported by the metabolome data described below. Two observations in the transketolase depleted strain suggested that the presence of ribose in growth medium could influence NADH/NADPH metabolism. One, rescue of the growth defect of Δ*tkt* Δ*udhA* strain in basal medium by glycerol and g*lpK*^C^ allele was observed only in the presence of ribose (Fig. 3C). Two, the growth defect of Δ*tkt* mutant in the presence of glucose, following *pntAB* over- expression, was rescued by ribose supplementation (Fig. 4C). We hypothesized, during growth of Δ*tkt* mutant in glucose, allowing accumulation of ribose-5-P lowered the metabolic flux through Oxi-PPP by inhibiting the activity of Zwf and/or Gnd enzymes and redirected flux through glycolysis. To test this, we asked if ribose-5-P could directly inhibit the activity of these enzymes. Addition of ribose-5-P to cell free extracts made from wild type strain, lowered the specific activity of Zwf and Gnd enzymes in a concentration dependent manner. The specific activity decreased 30% and 40% for Zwf and Gnd respectively, in the presence of 10 mM ribose-5-P, with greater than 90% of this inhibition achieved at 5 mM ribose-5-P (Fig. 5, Table S1). In cell free extracts, before the addition of ribose-5-P, ratio of specific activity of Zwf to Gnd was ∼0.9, comparable to that reported (Mcnair and Chu, 1959).

**Fig. 5.**
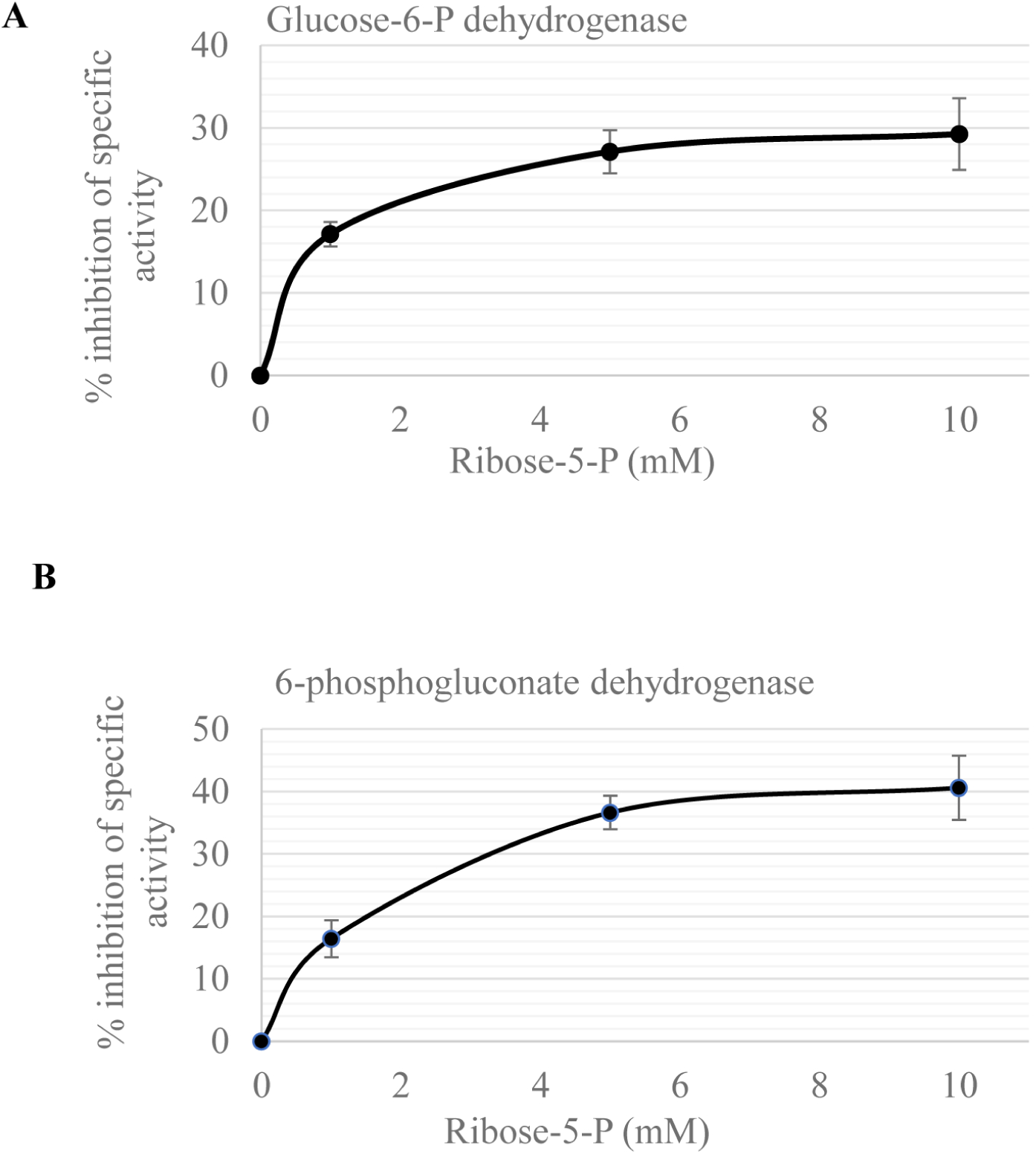
Inhibition of glucose-6-P dehydrogenase and 6-phosphogluconate dehydrogenase activity by ribose-5-P. The wild type strain (MG1655) was cultured to mid-log phase in LB, washed and re-suspended in Tris buffer. Following cell lysis, the enzyme activity was measured in the supernatant from three independent experiments as described under materials and methods. The error bars represent standard deviation of the mean.

The biochemical data described above can be used to explain the ribose sensitivity of Δ*tkt* mutant in basal medium. Under the gluconeogenic growth conditions, either the transketolase or the UdhA activity was required for growth (Fig. 3A) as one of the two activity is necessary for maintenance of sufficient NADH pool. Therefore, in the Δ*tkt* mutant, UdhA dependent synthesis of NADH from the oxidation of NADPH was essential for growth. Following accumulation of ribose-5-P, as the activity of Zwf and Gnd enzymes are inhibited, sufficient NADPH may not be available for UdhA mediated synthesis of NADH, leading to growth arrest. It other words, it can be said that, ribose sensitivity of Δ*tkt* mutant in basal medium was also consequence of NADH restriction and mechanistically similar to the Δ*tkt* Δ*udhA* growth defect. Therefore, the growth defect in both strains could be rescued by activating the glycolytic flux leading to GAPDH dependent NADH synthesis (Fig. 1 and Fig. 3).

### Transketolase function alleviates growth defect arising from the perturbation of NADH metabolism by GapN expression

The growth of wild type strain and Δ*tkt* mutant are comparable in basal medium containing glucose. However, when the *gapA* gene, encoding NAD^+^- dependent GAPDH was replaced with the *gapN* gene from *Streptococcus Mutans* (referred as Δ*gapA*::*gapN*) that codes for NADP^+^ - dependent GAPDH (Centeno-Leija *et al*., 2013), the growth of Δ*tkt* mutant was much more severely inhibited as compared to the wild type strain (Fig. 6A). Replacing *gapA* with *gapN* would result in the synthesis of two molecules of NADPH (instead of two molecules of NADH), for every molecule of glucose oxidized by GAPDH. This would be expected to lower the cellular NADH to NADPH ratio. To test if this perturbation was the cause of growth inhibition, UdhA, which catalyzes NADH synthesis concomitant with oxidation of NADPH was expressed from plasmid to counteract any reduction in NADH to NADPH ratio. Growth defect was rescued following the over-expression of *udhA* from plasmid (Fig. 6B), which indicated that the decrease in NADH to NADPH ratio was responsible for the weak and strong growth inhibition observed in the Δ*gapA*::*gapN* and Δ*tkt* Δ*gapA*::*gapN* strains respectively.

**Fig. 6.**
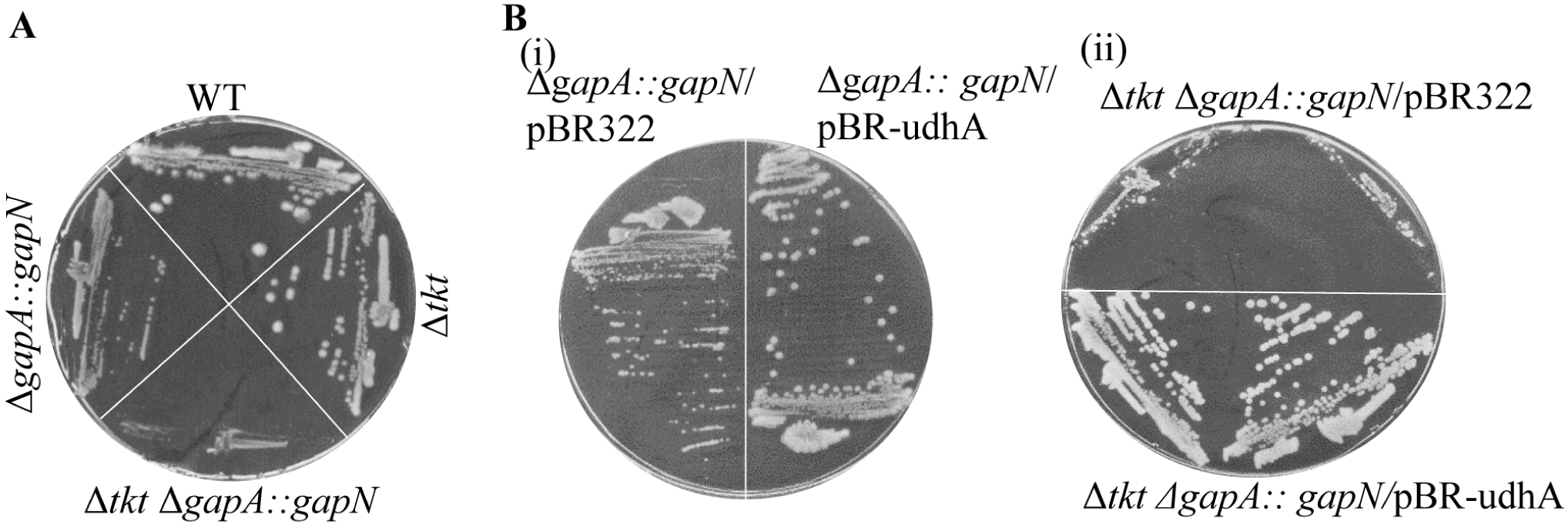
Over expression of UdhA rescued growth defect of strain carrying the Δ*gapA::gapN* allele. Strains whose relevant genotypes are indicated were streaked on basal medium supplemented with glucose (A, B). Antibiotics was added when strains carried plasmid. WT (MG1655), Δ*tkt/*pAM*-*tktB (AV104), *ΔgapA:: gapN* (AV245), Δ*tkt ΔgapA:: gapN/*pAM*-tktB* (AV246), *ΔgapA::gapN*/pBR322 (AV310), *ΔgapA::gapN*/pBR-udhA (AV311), Δ*tkt ΔgapA:: gapN/* pAM*-*tktB/pBR322 (AV314), Δ*tkt ΔgapA:: gapN/*pAM*-* tktB/pBR-udhA (AV315).

The more severe growth inhibition in Δ*tkt* Δ*gapA*::*gapN* strain (as compared to isogenic Δ*gapA*::*gapN* strain) suggested that transketolase activity could alleviate the decrease in NADH to NADPH ratio during growth on glucose. We propose, the decrease in NADH to NADPH ratio from the presence of Δ*gapA*::*gapN* allele could provide a signal for transketolase mediated conversion of the glycolytic intermediates, fructose-6-P and glyceraldehyde-3-P to xylulose-5-P and erythrose-4-P, thereby lowering glycolytic flux and GapN dependent NADPH synthesis.

The Δ*gapA*::*gapN* mutant did not grow in basal medium or minimal medium with glucose as the sole carbon source (data not shown). The absence of growth in basal medium can be attributed to the irreversible nature of reaction catalyzed by GapN, which is a non- phosphorylating type NADP^+^ dependent GAPDH uni-directionally producing 3- phosphoglycerate (Spaans *et al*., 2015) and hence cannot support gluconeogenesis. The reason for growth defect in medium containing glucose as the sole carbon source is unclear.

### Genetic changes that rescue growth defect of transketolase mutant in LB Medium

Growth of the Δ*tkt* /pAM-tktB strain was IPTG dependent – withdrawal of IPTG resulted in growth arrest in broth (Fig. 1A, i) and LB agar plates (Fig. 1A, ii). Addition of pyridoxine, shikimic acid and aromatic amino acids tyrosoine, phenyalanine and tryptophan did not rescue growth of Δ*tkt* mutant in the absence of IPTG (data not shown). To understand the molecular basis for growth defect of Δ*tkt* mutant in LB medium, transposon mediated mutagenesis was carried out as described under methods and transposon insertions that suppressed the growth defect were identified. Sequencing across the transposon-chromosome junction revealed five independent insertions in the *deoB* ORF, one in *deoD* and one at the end of the *glpF* gene, the first gene of the glp*FKX* operon. Two of the insertions in *deoB* and the insertion at the end of *glpF* were characterized and reported previously (Vimala and Harinarayanan, 2016). Using a multi-copy plasmid library of *E. coli* genes, we identified the *pntAB* genes as suppressor of the growth defect. Experiments to understand the basis for the suppression of growth defect by each genetic change is described below.

#### (i) ΔdeoB / ΔdeoD

The *deoB* and *deoD* genes code for phosphopentomutase and purine nucleoside phosphorylase respectively, and are part of the *deoCABD* operon. We expected transposon insertions in *deoB* or *deoD* would result in the inactivation of gene harbouring the insertion, and also that insertions in *deoB* could have a polar effect on the expression of *deoD*. Therefore, to test the role of each gene function in the growth defect of tranketolase depleted strain, we created non-polar Δ*deoB*::FRT or Δ*deoD*::FRT alleles from the Δ*deoB*::kan or Δ*deoD*::kan alleles available in the keio collection (Baba *et al*., 2006). Each allele suppressed the growth defect of the Δ*tkt* strain, indicating that loss of either the phophopentomutase (DeoB) or purine nucleoside phosphorylase (DeoD) could rescue the growth defect (Fig. 7B). DeoD and DeoB enzymes catalyze sequential reactions. DeoD cleaves purine ribonucleosides and purine deoxyribonucleosides into nulceobase and ribose-1-P or deoxyribose-1-P and DeoB (phosphopentomutase) converts ribose-1-P to ribose-5-P (Hammer-Jespersen and Munch- Petersem, 1970) Therefore, synthesis of ribose-5-P from nucleoside would be lowered in *deoD* mutant and completely blocked in the *deoB* mutant. This suggested, reduction in ribose-5-P synthesis from nucleosides present in the LB medium could be the reason for growth rescue by the mutations. Consistent with this idea, ribose supplementation to LB inhibited the growth of *Δtkt* Δ*deoB*::Kan/pAM-tktB strain and this was dependent on *rbsK* gene function required for the conversion of ribose into ribose-5-P (Fig. 7C). The supplementation of nucleosides inhibited growth of Δ*tkt*/pAM-tktB strain in basal minimal medium and this was rescued by *deoB* inactivation (Fig. S1A). These results indicated nucleoside degradation as an important source of ribose-5-P during growth in LB and its accumulation conferred growth inhibition in the transketolase deficient strain. From these observations, it can be said, (i) the molecular basis for growth inhibition of transketolase depleted strain in LB medium and ribose sensitivity of the strain in defined medium are the same and arises from the accumulation of ribose-5-P (and possibly other pentose phosphate) (ii) nucleosides serve as a carbon source during growth in LB through the generation of ribose-5-P.

**Fig. 7.**
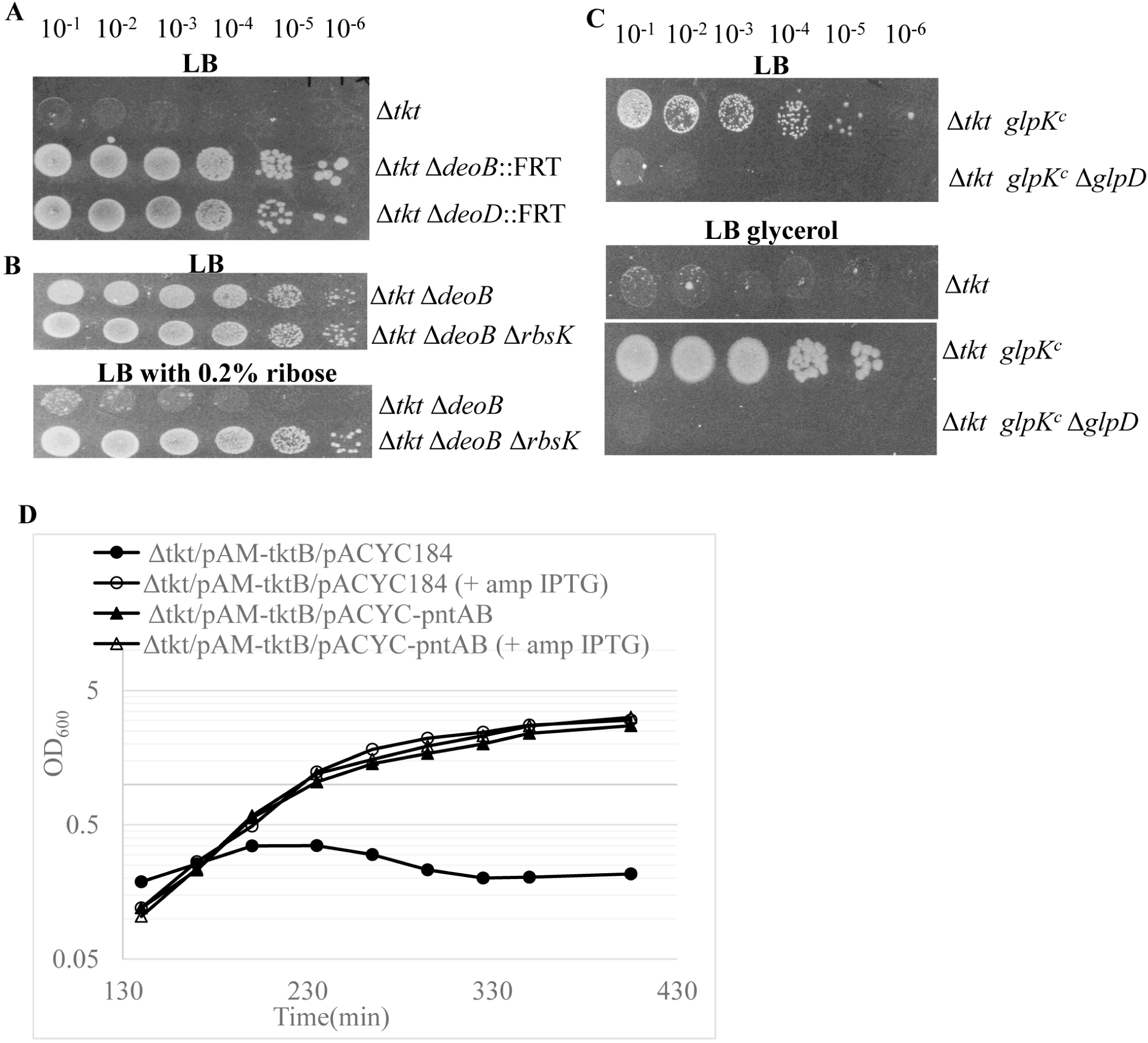
Growth defect of Δ*tkt* mutant in LB and its rescue by loss of *deoB or rbsk* function, activation of glycerol metabolism or increased expression of pntAB. Strains whose relevant genotypes are indicated were depleted for transketolase activity, serially diluted and spotted on LB plates and LB plate with the indicated supplements (A - C). Δtkt/pAM-tktB/pACYC184 and Δtkt/pAM-tktB/pACYC-pntAB strains cultured overnight in LB cm amp IPTG were washed and sub-cultured into LB medium with chloramphenicol and with (○, Δ) or without (● ,▴) ampicillin and IPTG, and the increased in A600 was monitored (D). Strains used are Δ*tkt*/pAM-tktB (AV104); Δ*tkt deoB*::FRT/pAM*-*tktB (AV159); Δ*tkt deoD*::FRT/pAM*-*tktB (AV135); Δ*tkt deoB/*pAM*-*tktB (AV155); Δ*tkt deoB rbsK/* pAM*-*tktB (AV160) ; Δ*tkt glpK^c^/* pAM*-*tktB (AV148);Δ*tkt glpK^c^ glpD/*pAM*-*tktB (AV150) ; Δ*tkt/* pAM*-*tktB/pACYC184 (AV110) and Δ*tkt/*pAM-tktB/pACYC-pntAB (AV112) .

#### _(ii)_ *glpK^C^*

The transposon insertion at the end of the *glpF* gene (coding for glycerol permease) increased the expression of the downstream *glpK* gene (coding for glycerol kinase) and made it constitutive (Vimala and Harinarayanan, 2016). *glpK*^C^ rescued the growth of Δ*tkt* strain in LB medium, and this was further enhanced by glycerol supplementation (Fig. 7C), suggesting that, strength of growth rescue was proportionate to glycerol catabolism. Catabolism of the endogenously synthesized glycerol (Weichart et al. 1993) may explain the growth rescue by *glpK*^C^ in LB. Glycerol supplementation did not support growth of Δ*tkt* strain in LB medium (Fig. 7C, third row), which indicated, *glpK^C^* allele was necessary for glycerol catabolism in the absence of transketolase activity. This is consistent with our finding that the *glpFKX* promoter can be repressed during the accumulation of ribose-5-P and possibly other pentose phosphates (Vimala and Harinarayanan, 2016).

To further characterize the basis for rescue of Δ*tkt* growth defect by *glpK*^C^, the role of glycerol- 3-P dehydrogenase and triose phosphate isomerase activity in the rescue was examined. Introduction of Δ*glpD*::kan (Fig. 7C) or the ΔtpiA::kan (Fig. S1B) mutation into the Δtkt *glpK*^C^ strain resulted in loss of the growth rescue with or without glycerol. Inactivation of glycerol kinase, using the ΔglpK::kan also resulted in loss of growth rescue (data not shown). Together, these results suggested, the biosynthesis of glycolytic intermediate glyceraldehyde-3-P from glycerol was necessary for suppression of growth defect. Thus, as seen for rescue of ribose sensitivity of Δ*tkt* strain in basal medium by glycerol (Fig. 1D), synthesis of glyceraldehyde- 3-P from glycerol was required for growth rescue in LB medium.

#### (iii) Multi-copy pntAB

Over-expression suppressors that rescued the growth defect of Δ*tkt* strain in LB were identified by transforming *E. coli* gene library constructed in plasmid pACYC184 into the Δ*tkt*/pAM- tktB strain and screening for IPTG independent growth. As would be expected, four independent plasmid clones with DNA fragments of different length and carrying the *tktA* gene were obtained. In addition, one plasmid clone carrying the *pntAB* genes that code for the membrane bound proton translocating pyridine nucleotide transhydrogenase was obtained (pACYC-pntAB). Introduction of pACYC-*pntAB* into the Δ*tkt*/pAM-tktB strain supported IPTG-independent growth (Fig. 7D). Since studies have suggested PntAB to primarily function in the synthesis of NADPH by the oxidation of NADH (Sauer *et al*., 2004, Spielmann *et al*. 2018), it was possible, suppression of growth defect by multi-copy *pntAB* could follow from an increase in the cellular NADPH pool. Results described below are also consistent with this idea, and importantly, growth conditions and genetic changes that rescued the growth defect of Δ*tkt* strain were dependent on *pntA* gene function. Multi-copy *pntAB* did not rescue the ribose sensitivity of Δ*tkt* strain in basal medium (data not shown).

### Growth defect of transketolase depleted strain in LB was rescued by glucose, gluconate or the Δ*gapA*::*gapN* allele

The rescue of growth defect of transketolase depleted strain in LB medium and LB containing ribose by loss of DeoB or DeoB and RbsK functions respectively (Figs. 7B) indicated the growth defect in LB and ribose sensitivity in defined medium (Fig. 1B) are mechanistically related. Since glucose or gluconate supplementation rescued ribose sensitivity of transketolase depeleted strain (Fig. 1C and 1E), we tested their ability to rescue the growth defect in LB medium. Glucose supplementation rescued the growth defect and this was dependent on Pgi and Zwf functions (Fig. 8A rows 2 and 3), indicating, glucose catabolism through glycolysis and as well as Oxi-PPP pathway was required for growth rescue. Gluconate rescued the growth defect of Δ*tkt* mutant and as would be expected, the rescue was dependent on Edd function (Fig. 8B). As pyruvate is produced during glucose and gluconate catabolism, we tested pyruvate, but it did not rescue the growth defect (data not shown). Together, these results suggested, as observed for ribose sensitivity of transketolase depleted strain in defined medium, increase in the glycolytic flux between glyceraldehyde-3-P and pyruvate may be important for growth rescue.

**Fig. 8.**
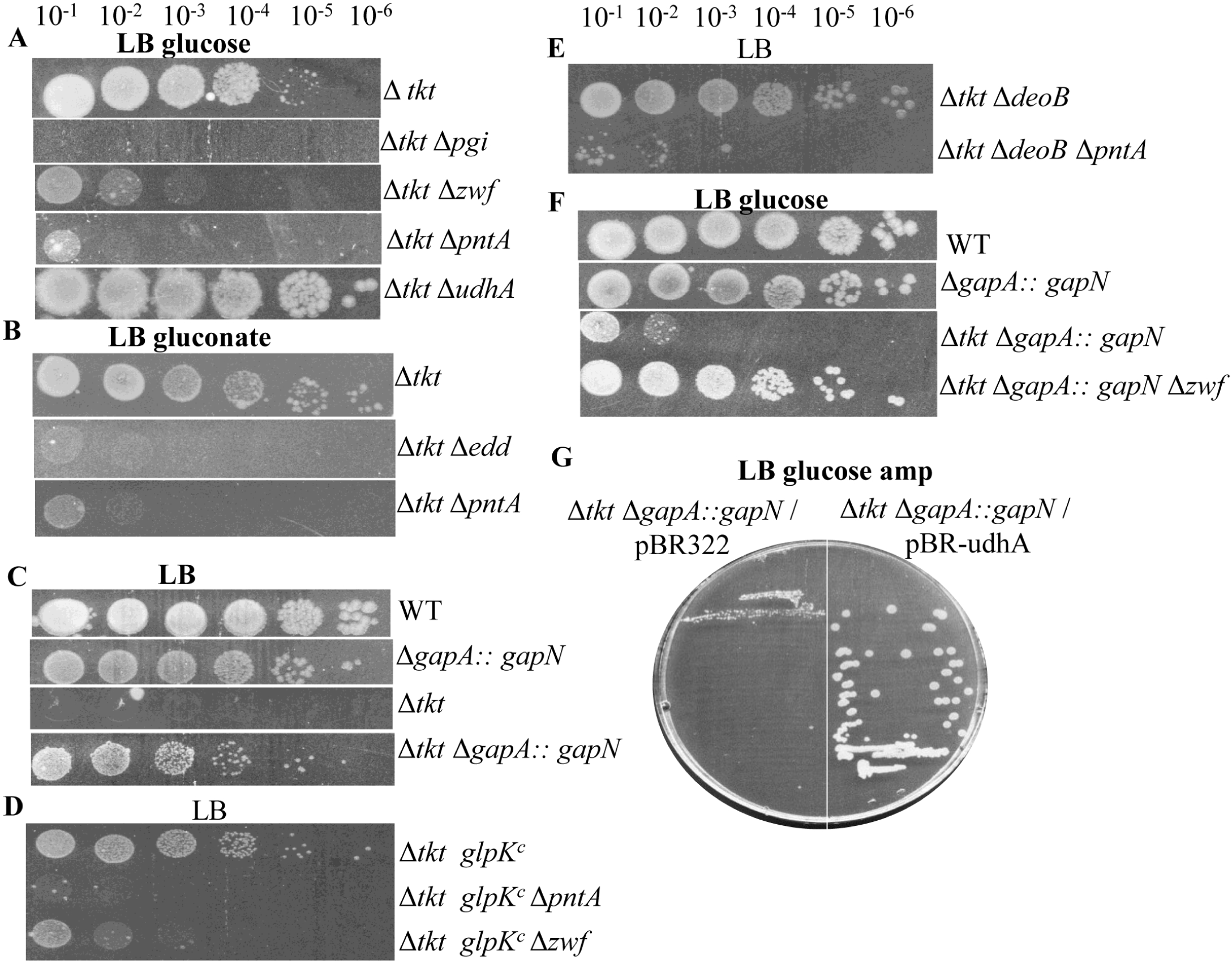
Genetic changes that perturb NADPH pool affect the growth of the transketolase mutant. Strains whose relevant genotypes are indicated were depleted for transketolase activity, serially diluted and spotted (A – F) or streaked (G) on LB plate with the sugar supplements where indicated. WT(MG1655); Δ*tkt*/pAM*-*tktB (AV104); Δtkt Δ*pgi*/pAM*-*tktB (AV108); Δ*tkt* Δ*zwf*/pAM*-*tktB (AV107); *Δtkt Δedd*/pAM*-*tktB (AV109); Δ*tkt* Δ*pntA*/pAM*-* tktB (AV130); Δ*tkt* Δ*udhA*/pAM*-*tktB (AV208);; *ΔgapA:: gapN* (AV245); Δ*tkt ΔgapA:: gapN*/pAM*-*tktB (AV246); Δ*tkt glpK^c^*/pAM*-*tktB (AV148); Δ*tkt glpK^c^* Δ*pntA*/pAM*-*tktB (AV151); Δ*tkt glpK^c^* Δ*zwf*/pAM*-*tktB (AV149); Δ*tkt* Δ*deoB*/pAM*-*tktB (AV155); Δ*tkt* Δ*deoB* Δ*pntA*/pAM*-*tktB (AV186); Δ*tkt ΔgapA:: gapN* Δ*zwf*/pAM*-*tktB (AV250); *Δtkt ΔgapA::gapN*/pAM*-*tktB/pBR322 (AV314) and *Δtkt ΔgapA::gapN*/pAM*-*tktB/pBR-*udhA* (AV315).

To study the role, in any, for GAPDH dependent NADH synthesis in the growth rescue, the NADH generating GapA function was replaced with the NADPH generating GapN by introducing the ΔgapA::gapN allele. The ΔgapA::gapN allele by itself did affect growth significantly, but the growth defect of transketolase depleted strain was rescued by the ΔgapA::gapN allele (Fig. 8C). This showed, NADH synthesis by GapA was not necessary for growth rescue and that an increase in the NADPH pool may be an important requirement to rescue growth defect of transketolase deficient strain in LB. A similar experiment to address the role of NADPH in the ribose sensitivity phenotype of transketolase deficient strain could not be performed, because the ΔgapA::gapN strain does not grow in basal medium.

### Growth defect from transketolase depletion in LB medium arises from NADPH limitation

Rescue of the growth defect of transketolase deficient in LB by, (i) genetic changes that lower cellular ribose-5-P pool (Fig. 7B); (ii) multi-copy *pntAB* (Fig. 7D) and (iii) presence of Δ*gapA*::*gapN* allele (Fig. 8C), taken together with the finding that ribose-5-P inhibited the activity of NADPH generating Zwf and Gnd enzymes (Fig. 5), suggested the growth defect may be related to NADPH limitation. The results described below lend further support to this idea. Growth rescue by *glpK*^c^ allele was dependent on Zwf or PntA functions, each of which supports NADPH synthesis (Fig. 8D). Growth rescue by Δ*deoB* allele was dependent on PntA function (Fig. 8E). The Δ*pntA* or Δ*zwf* mutations individually do not perturb growth of Δ*deoB* (*tkt*^+^) or glpK^c^ (*tkt*^+^) strains in LB medium, indicating the requirement of Pnt or Zwf function for growth was specific to the transketolase deficient strain. The rescue of growth defect by glucose was dependent on *pntA* gene function and enhanced by the loss of *udhA* gene function (Fig. 8A rows 4 & 5). As PntA contributes to NADPH synthesis and UdhA lowers NADPH pool during growth on glucose (Canonaco, et al. 2001; Sauer *et al*., 2004), the results suggest, strength of the growth rescue by glucose may be proportionate to cellular NADPH pool. The growth rescue by gluconate was dependent on PntA function (Fig. 8B row 3).

We also found evidence that excess NADPH synthesis could be detrimental to growth of Δ*tkt* mutant in LB. Growth rescue of transketolase depleted strain by Δ*gapA*::*gapN* allele was impaired in the presence of glucose, where NADPH synthesis can be expected from both glycolytic and Oxi-PPP mediated catabolism of glucose. Finally, we observed, loss of Zwf function or over-expression of UdhA, each of which would lower the cellular NADPH pool, rescued growth defect of transketolase depleted strain (Figs. 8F & 8G). Overall, these results indicate, transketolase activity enabled the cell to better tolerate fluctuation (both decrease and increase) in the NADPH pool.

### Depletion of transketolase activity led to drop in pyridine cofactor pool and was alleviated under conditions that rescued the growth defect

Since the genetic data suggested growth defect of Δ*tkt* mutant in LB was associated with perturbation of NADPH pool, we tested this directly. The concentration of pyridine cofactors was measured in the wild type strain MG1655 and in the Δ*tkt*/pAM-tktB strain before and after depletion of transketolase activity. For carrying out depletion, overnight cultures grown in the presence of IPTG were washed, diluted and grown in the absence of IPTG until growth arrest (as described in methods). The concentration of pyridine nucleotide co-factors was compared between strains after normalizing the optical density of the cultures.

The concentration of each pyridine nucleotide co-factor, namely, NAD^+^, NADH, NADP^+^ and NADPH was measured using the cycling assay (Zhou et al. 2011). The NAD^+^ to NADH ratio was 9 and 12 respectively, in the wild type and the Δtkt/pAM-tktB strain without transketolase depletion (Fig. 9A). The NADP^+^ to NADPH ratio was 1.8 and 2.1 respectively in the wild type and Δtkt/pAM-tktB strain without transketolase depletion (Fig. 9B). Following depletion of transketolase activity and growth arrest, the NAD^+^ pool decreased ∼8 -fold, NADP^+^ dropped ∼ 2-fold as compared to the un-depleted strain; NADH and NADPH levels were below the detection limits of the assay (Figs. 9A and 9B).

**Fig. 9.**
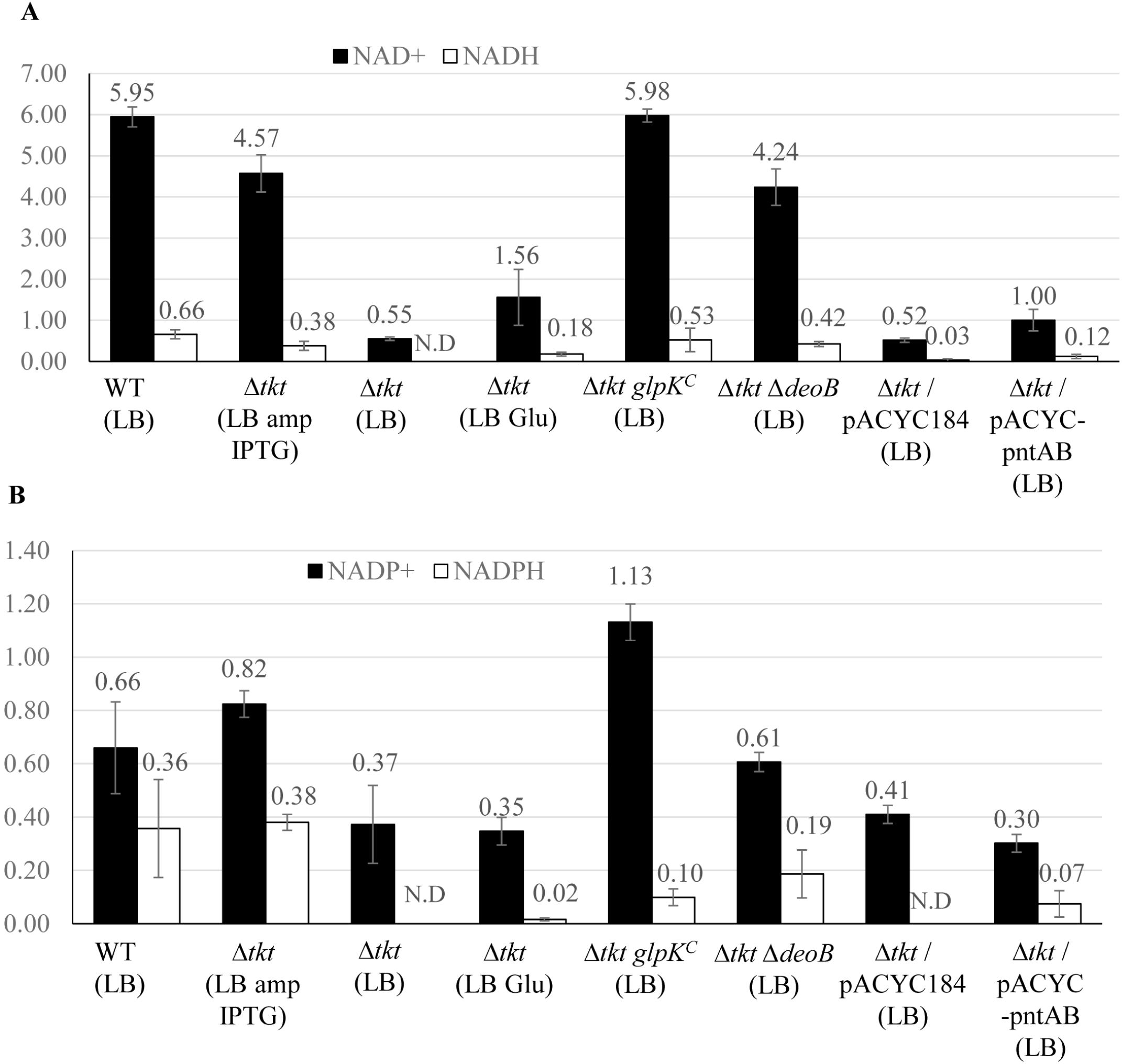
Effect of transketolase depletion on pyridine cofactor concentration. The transketolase activity in strains whose relevant genotypes are indicated were depleted using the IPTG dependent replication plasmid pAM-tktB and pyridine cofactor pools were measured in cultures normalized for A600 using the cycling assay (see methods for details). The assays were carried out using three independent cultures for each strain and values reported are in (µmoles/ gram dry weight). WT (MG1655); Δ*tkt*/pAM*-*tktB (AV104); Δ*tkt glpK^C^*/pAM*-*tktB (AV148); *Δtkt ΔdeoB*/pAM*-*tktB (AV155); Δ*tkt*/pAM*-*tktB/pACYC184 (AV110) and Δ*tkt*/pAM*-*tktB/pACYC-pntAB (AV112). N.D - Not Detectable

To find out if changes in the co-factor levels was specific to depletion of transketolase activity or a feature associated with growth arrest, we induced growth arrest by depletion of the essential protein Rho involved in transcription termination (Ref) using the conditional replication plasmid system employed for transketolase depletion. Following depletion of Rho activity and growth arrest, NAD^+^ and NADH level decreased ∼2-fold, NADP^+^ increased slightly, and NADPH dropped 4-fold, as compared to the un-depleted culture (Fig. S2). As changes in pyridine cofactor levels after transketolase depletion were more severe than that seen following Rho depletion, the results support the idea that depletion of transketolase function causes a specific pronounced decrease in pyridine cofactor pools over and above that associated with growth arrest.

Pyridine cofactor levels was measured following growth in the presence of glucose or genetic changes (deoB, glpK^C^ and multi-copy pntAB) that rescued growth defect of transketolase depleted strain. The deoB and glpK^C^ mutations showed robust rescue of cofactors, whose concentration were similar to that seen in the un-depleted strain; only weak rescue was observed with glucose and pACYC-pntAB. These results together with the genetic data lend support the idea that transketolase activity was required for maintenance of pyridine cofactor pool during growth in LB and probably other growth conditions that support synthesis of ribose-5-P and/or other pentose phosphates.

### Metabolome changes associated with depletion of transketolase activity

The metabolome of the strain subjected to transketolase depletion was compared with the metabolome of the strain without the depletion and as well as the wild type strain (all strains were grown in LB medium) by GC/MS-TOF (Gas Chromotography-Time-of-Flight/Mass spectrometry) using two biological replicates for each strain (Table S4), see methods for details. The mean value was used for calculating fold-change in concentration of metabolites between the strains. Identifiable metabolites whose levels were altered 1.5-fold or more in the transketolase depleted strain as compared to the strain not depleted for transketolase activity or the wild type strain were identified (Fig. 10). In agreement with the genetic data that pentose phosphates can accumulate in transketolase deficient strain, the metabolomics data showed ribulose-5-P level to be elevated ∼9- and ∼19-fold in the transketolase depleted strain as compared to non-depleted and wild type strain respectively. However, it is not clear if GC/MS-TOF would distinguish the major pentose phosphates that have identical molecular weight (ribose-5-P, ribulose-5-P, xylulose-5-P and arabinose-5-P), therefore, it may be also possible, the data could represent the sum total of all pentose phosphates. Ribose was elevated 5- and 2.25-fold and ribitol 3.9- and 4.8-fold in the non-depleted and wild type strain respectively as compared to the transketolase depleted strain. It is not clear if these are compounds are naturally produced in the *E. coli* metabolome or produced under the experimental conditions. Consistent with the genetic data of decrease in glycolytic flux after transketolase depletion, the glycolytic intermediate, 3-phosphoglycerate was 1.75 and 2.5-fold lower in the transketolase depleted strain as compared to the non-depleted and wild type strain respectively. TCA cycle intermediates, citric acid, alpha ketoglutaric acid, succinic acid and nucleotides, AMP, UMP and TMP were significantly lowered in the transketolase depleted strain as compared to the non-depleted and wild type strain. Nicotinamide, a product of the NAD^+^ salvage pathway (Gholson *et al*., 1969) was ∼ 3-fold lowered in the transketolase deficient strain, this may be expected since NAD^+^ pool was lowered ∼10-fold in the transketolase depleted strain (Fig. 10). The glutathione pool was 4-and -6 fold lower in the transketolase depleted strain as compared to the non-depleted and wild type strain respectively. It was possible, this may be related to the decrease in NADPH pool in the transketolase depleted strain (Fig. 10) and consequent loss of glutathione regeneration from glutathione disulphide by glutathione reductase which uses NADPH as cofactor. The decreased glutathione pool in the transketolase depleted strain also corroborates with the significant drop in cysteine-glycine dipeptide, which is the product of glutathione hydrolysis by glutathione hydrolase proenzyme gamma glutamyl hydrolase, Ggh (Suzuki et al.1986a, Suzuki et al. 1986b).

**Fig. 10.**
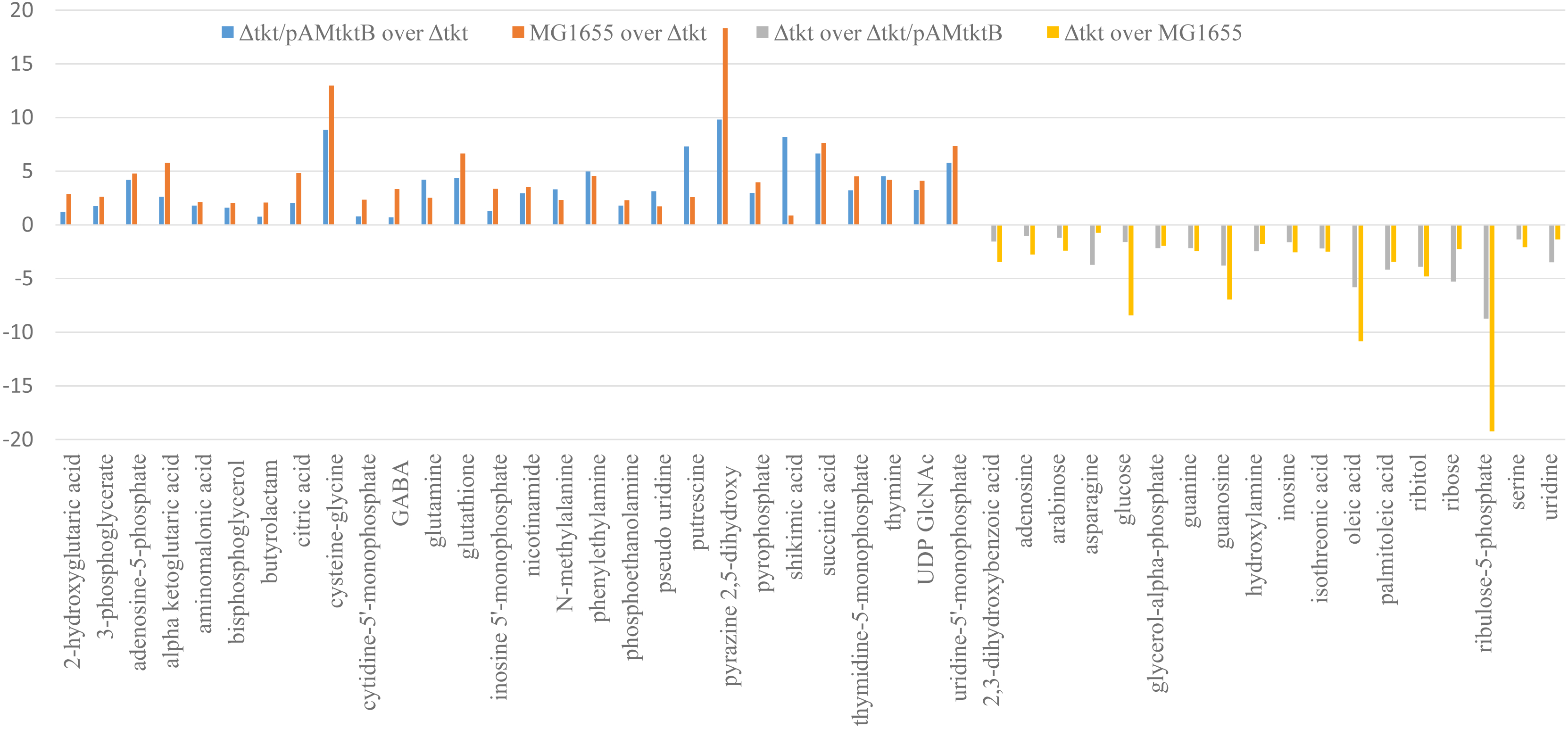
List of metabolites whose levels were altered 1.5- fold or greater following depletion of transketolase activity. The positive and negative values indicate higher and lower concentration respectively of metabolites in the wild type and strain without depletion of transketolase activity as compared to isogenic strain depleted for transketolase activity. The strains were cultured in LB medium and values reported are the average from two independent experiments.

## Discussion

In this study, evidence was presented for role of transketolase function in modulation of NADPH/NADH metabolism in *Escherichia coli*, which to the best of our knowledge may be the first report. Since transketolase function is conserved across bacteria, these findings could be relevant to the physiology of other bacteria as well. Biochemical assays showed that NAD^+^, NADH and NADPH pools were significantly lowered in transketolase deficient strain under growth conditions that would lead to the accumulation ribose-5-P and other pentose phosphates. The genetic data indicated, this was primarily brought about by the regulation of the metabolic flux through the central carbon metabolic pathways of glycolysis and Oxi-PPP. As discussed below, in growth medium containing casamino acids as sole carbon source, where metabolism is primarily gluconeogenic, transketolase function was required for maintenance of optimal NADH pool, while in LB medium it supported the maintenance of NADPH pool by preventing the accumulation of ribose-5-P (and other pentose phosphates) that inhibited Zwf and Gnd enzyme activity. The role of transketolase function in the maintenance of pyridine nucleotide pool is discussed below for each of the following growth condition, (i) defined medium with casaminoacids as primary carbon source (gluconeogenesis) (ii) defined medium with casaminoacids and glucose (glycolysis) and (iii) LB medium.

### Role of transketolase in regulation of NADH pool under gluconeogenic growth conditions

During growth with amino acids as the sole carbon source, the transketolase depleted strain exhibited two phenotypes, namely, synthetic growth defect with loss of UdhA function and growth inhibition by ribose (ribose sensitivity). The relationship between the two phenotypes are discussed below.

Since UdhA transhydrogenase catalysed the reduction of NAD^+^ concomitant with oxidation of NADPH (Canonaco, et al. 2001, Sauer, *et al*. 2004), a reduction in NADH to NADPH ratio may be expected in the Δ*udhA* mutant. The synthetic growth defect of Δ*udhA* Δ*tkt* strain (Fig. 3A) suggested the imbalance was further enhanced by the loss of transketolase function. We propose, during gluconeogenic growth, transketolase function supported GAPDH dependent NADH synthesis by catalysing synthesis of fructose-6-P and glyceraldehyde-3-P (from pentose phosphates) and increasing the glycolytic flux. Support for this idea comes from the rescue of Δ*udhA* Δ*tkt* growth defect by glucose or gluconate (Fig. 3B), sugars that are catabolized through glycolysis and support GAPDH dependent NADH synthesis.

Ribose inhibited growth of transketolase depleted strain in casamino acids medium (Fig. 1B). Accumulation of ribose-5-P was expected due to inactivation of non-Oxi-PPP pathway. Since accumulation of ribose-5-P conferred growth inhibition in the transketolase depleted strain in LB (Figs. 7A and 7B) we think the same can explain ribose sensitivity in basal medium. Since Zwf and Gnd enzyme activities were progressively inhibited by an increase in concentration of ribose-5-P (Fig. 5), this would suggest reduced flux through the Oxi-PPP pathway and lower NADPH pool in the transketolase depleted strain. Measurements showed NADPH pool to be below detection limit (Fig. 10B).

Sugars that are catabolized through glycolysis rescued the ribose sensitivity of transketolase depleted strain (Fig. 1B to 1E) and as well as the Δ*tkt ΔudhA* growth defect (Fig. 3A & 3B). This suggested a common suppression mechanism for the two growth defects. Since glycolysis is associated with NADH synthesis, an increase in the NADH level was the likely reason for rescue in both scenarios. How can the ribose sensitivity of transketolase depleted strain, which our data indicate to arise from the inhibition of Zwf and Gnd activities and the consequent decrease in NADPH pool be rescued by an increase in the NADH level? We propose, the drop in NADPH pool impaired UdhA mediated synthesis of NADH, and ultimately this was the cause of growth inhibition in the transketolase depleted strain. It can be said, the ribose sensitivity of Δ*tkt* mutant and growth defect of Δ*tkt ΔudhA* strain arises from NADH depletion.

### Role of transketolase in regulation of NADH pool during glucose catabolism

The physiological significance of transketolase function during glucose catabolism, as deduced from the growth properties of transketolse depleted strain in media with glucose and casaminoacids is discussed below. The significance of maintaining transketolase dependent metabolic flux into glycolysis was revealed by the growth defect of Δ*pgi* Δ*tkt* strain in basal medium containing glucose and its rescue by gluconate or glycerol (in the presence of *glpK*^C^ allele), but not pyruvate (Fig. 2). This suggested, an increase in the glycolytic flux between glyceraldehyde-3-P and pyruvate could be the likely cause for rescue of growth defect. Since this section of the glycolytic pathway contributes to GAPDH dependent NADH synthesis, and evidence from this study point to perturbation of pyridine cofactor pool in the transketolase depleted strain, we favour NADH depletion as the likely cause for growth defect of Δ*pgi* Δ*tkt* strain. The role of transketolase function in maintenance of NADH pool during growth on glucose was supported by two other observations, (i) reduced tolerance of Δ*tkt* mutant (relative to wild type strain) to increased expression of PntAB transhydrogense (Fig. 4A); (ii) reduced tolerance of Δtkt mutant to the expression of GapN (in place of GapA) (Fig. 6A) both of which would lower the NADH and increase the NADPH pool in the cell. These results suggest, transketolase activity is required during growth conditions that lower NADH to NADPH ratio. Rescue of the growth defect due to GapN expression by the over-expression of UdhA lends further support this idea (Fig. 6B, ii). A previous study reported that replacing the *gapA* gene with *gapN* reduced growth rate of *E. coli* in glucose due to the poor catabolism of glucose (Centeno-Leija *et al*., 2013). Since UdhA over-expression rescued the growth defect of such a strain (Fig. 6B, i), we suggest, NADH limitation was responsible for the poor growth of the *gapN* strain in glucose.

### Role of transketolase in the regulation of NADPH pool during growth in LB

LB being a complex undefined medium, the nature of metabolic flux during growth in LB is ill defined. However, comparing growth properties of transketolase depleted strain in LB medium and defined medium suggested the metabolic flux regulation by transketolase and consequent effect on NADH/NADPH metabolism are different in the two growth conditions, as discussed below.

Our data strongly suggest, the growth defect of Δ*tkt* mutant in basal medium containing ribose and in the LB medium resulted from accumulation of ribose-5-P and the inhibition of Zwf and Gnd activities by ribose-5-P (Figs. 1B, 5, 7A&B). While the Oxi-PPP reactions directly support NADPH synthesis, it is also required for directing the metabolic flux into glycolysis through the non-Oxi PPP pathway and contribute to NADH synthesis.

In LB medium, the growth defect of transketolase deficient strain was rescued by (i) the Δ*deoB* or *glpK*^C^ mutation (Fig. 7B & 7C) (ii) increased expression of PntAB (Fig. 7D) and (iii) glucose or gluconate supplementation (Fig. 8A & 8B). Importantly, growth rescue by glucose and gluconate was dependent on PntA function and the rescue by glucose significantly improved following elimination of UdhA activity (Fig. 8A & 8B). However, none of the genetic changes or growth conditions that rescued growth defect of Δ*tkt* mutant in defined media showed dependence on PntA function. These observations strongly support NADPH limitation as the cause of growth defect in LB and NADH limitation the cause of growth inhibition in defined media.

In conclusion, results from this study have revealed a novel role for transketolase activity in *E. coli* physiology and possibly other bacteria. By regulating the metabolic flux between glycolysis and the pentose phosphate pathway, especially in growth conditions that support synthesis of pentose phosphates, transketolase function regulates the cellular NADH and NADPH metabolism.

### Experimental Procedures

#### Media and growth conditions

LB medium was prepared by hydration of a premade mix (0.5% yeast extract, 1% tryptone and 1% NaCl) from HIMEDIA (Cat. No. M1245). Minimal A salts medium was made as previously described (Miller, 1992). Solid media was prepared by adding agar (Difco) to a final concentration of 1.5% and sterilized by autoclaving. Plates and cultures were supplemented to the following final concentrations, glucose (0.2%), glycerol (0.2%), gluconate (0.2%), ribose (0.2%), casamino acids (0.2%), tryptophan (100 µg/mL), shikimic acid (100 µg/ml), nucleosides (50 µg/mL each of adenosine, guanosine, cytosine, uridine and thymidine) and pyridoxine (10 µM). Final concentration of antibiotics used, ampicillin (100µg/mL), kanamycin (25µg/mL), chloramphenicol (15µg/mL), tetracycline (10µg/mL) and spectinomycin (25µg/ml). Isopropyl-β-D-thiogalactopyranoside (IPTG) was used at a final concentration of 1mM unless mentioned otherwise. Cells were cultured in petri plates or in conical flasks with aeration (with media at one-tenth volume) using rotary shaker at 37°C. Strains with the Δ*tkt* mutation (ΔtktA::FRT ΔtktB::FRT) were maintained with plasmid pAM- tktB. Overnight cultures grown in LB medium with ampicillin and IPTG were sub-cultured at 1:1500 dilution into fresh LB medium (without ampicillin and IPTG) and used for dilution-spotting assays. After sub-culture, strains were grown to an optical density of 0.3 to 0.6 (A_600_) and cells equivalent to 1 O.D. was collected, washed and re-suspended in 1ml of minimal A salts, and serially diluted. Dilutions were spotted on appropriate plates and incubated at 37°C for up to 48 hours.

#### Bacterial strains, plasmid constructions and genetic procedures

All strains in the study are derived from the wild type *Escherichia coli* K-12 strain MG1655 and are listed in Table S1. Plasmids and Primers used are listed in Table S2 and Table S3 respectively. Standard procedures were followed for cloning into plasmids and transformation (Russell and Sambrook, 2001). Plasmid pAM-tktB was made by sub-cloning the *tktB* gene from pHR30 (Harinarayanan *et al*., 2008) into the *Eco*RI and *Hin*dIII sites of pAM34 (Gil and Bouché, 1991). Phage P1vir was used for transductions as described (Miller, 1992). *udh*A gene together with its promoter was cloned into the *Hin*dIII and *Bam*HI sites of pBR322 after PCR amplification using primers JGMudhAFP1 and JGMudhARP1. A sp^R^ derivative of pAM-tktB was made by replacing the *bla* gene with *aadA* by recombineering using primers JGOspecFP and JGOspecRP and published protocols (Yu *et al*., 2000). When required, the kan^R^ marker was excised from the knock out alleles in keio collection using plasmid pCP20 for expression of FLP recombinase (Cherepanov and Wackernagel, 1995).

Transpositions were carried out using modified lambda phage 1324 that carries a derivative of transposon Tn10 with *cat* gene as described in Miller (1992). The position of the transposon in the genome was determined by RATE (random amplification of transposon ends) PCR (Ducey and Dyer, 2002) using primer JGO1324pcr that binds at the end of the transposon followed by sequencing using a nested primer JGO1324seq. Three insertions were mapped to the 1224 base pair *deoB* ORF at 89, 728, and 341 bases from the start site of the ORF. One insertion was mapped to *deoD* ORF and was 325 bases from start site of the ORF. To identify multi-copy suppressors of growth defect of Δ*tkt* strain, an *E. coli* genomic library constructed in the plasmid pACYC184 was transformed into the Δ*tkt* strain carrying plasmid pAM-tktB (AV94). Fast growing colonies were screened amongst Cm^r^ transformants obtained in the absence of IPTG. Plasmid clones isolated from fast growing colonies were sequenced using primers JGHtetA and JGHtetB.

#### Chemicals

All chemicals were obtained from Sigma Chemical Co., St Louis, Mo. β-Nicotinamide adenine dinucleotide (Cat. No. N8285), β-Nicotinamide adenine dinucleotide reduced disodium hydrate (Cat. No. N6660), β-Nicotinamide adenine dinucleotide phosphate sodium salt (Cat. No. N8035), β-Nicotinamide adenine dinucleotide 2-phosphate reduced tetrasodium salt (Cat. No. N9910), Thiazolyl blue tetrazolium bromide (Prod. No. M2128), Phenazine ethosulphate (Cat. No. P4544), Glucose-6-phosphate Dehydrogenase from baker’s yeast (*S. cerevisiae*) (Cat. No. G4134), Alcohol Dehydrogenase from *S. cerevisiae* (Cat. No. A3263), Glucose-6- phosphate sodium salt (Cat. No. G7879), D-Ribose 5-phosphate disodium salt dihydrate (Cat. No 83875) and 6-phosphogluconic acid trisodium salt (Cat. No. P7877).

#### Measurement of pyridine nucleotide cofactors

Cells equivalent to 0.6 OD (A_600_) was used for extraction of cofactors. For depletion of transketolase activity, the Δ*tkt /* pAM-tktB strain was cultured in LB medium containing ampicillin and IPTG and diluted 1:1500 into LB medium for depletion of transketolase activity. The MG1655Δ*rho/*pAM-rho strain was grown in LB medium containing ampicillin and IPTG and diluted 1:50000 into LB medium to get growth arrest from depletion of Rho activity. All assays were carried out with three independent cultures for each strain.

Cells were centrifuged at 15,000 rpm for 1 min and pellets were re-suspended in 300 *µ*l of 0.2M HCl (for NAD^+^, NADP^+^ extraction) or 0.2M NaOH (for NADH, NADPH extraction). The samples were placed in 50°C water bath for 10 min and then left on ice. The extracts were neutralized by adding 300 *µ*l of 0.1M NaOH (for NAD^+^, NADP^+^ extraction) or 300 *µ*l of 0.1MHCl (for NADH, NADPH extraction) added drop wise with vortexing. The cellular debris was removed by centrifuging at 15,000 rpm for 5 minutes and supernatant transferred to fresh tubes and stored at −20°C for less than 24 h before the assay.

The intracellular concentrations of pyridine nucleotide cofactors were measured using a sensitive cycling assay with minor modification (Zhou et al. 2011). For each assay, a reagent mixture consisting of 25µl each of 1.0 M Tris buffer (pH 8.0), absolute ethanol, 40mM EDTA (pH 8.0), 4.2mM MTT (3-[4,5-dimethylthiazol-2-yl]-2,5-diphenyltetrazolium bromide; thiazolyl blue), and 50µl of 16.6 mM PES (phenazine ethosulfate) was prepared. The following volumes were added to the wells of 96-well microtiter plate: 12.5µl neutralized cell extract, 75 µl water, and 150 µl reagent mixture and the mix was heated to 30°C. For measurement of NADH and NAD^+^, the reaction was started by adding 12.5µl (6.25U/reaction) of yeast ADH (340 U) made in 0.1 M Tris buffer (pH 8.0). The absorbance at 570 nm was recorded for 10 min at 30°C using the Varioskan Flash plate reader. The assay was calibrated using 2–16µM standard solutions of NAD+ and 0.5-6 µM standard solutions of NADH.

The intracellular NADPH and NADP^+^ concentrations were measured using a reagent mix with 25 µl of 10mM glucose-6-P (instead of ethanol) and the reaction was started by adding 12.5µl (0.342U//reaction) of baker’s yeast G6PDH (137 U) made in 5mM sodium citrate (pH 7.4). The assay was calibrated using 0.5-6 µM standard solutions of NADP+ and 0.5-5 µM standard solutions of NADPH.

The slope (ΔA/min) of the linear region of the absorbance versus time plot was used to calculate the concentration of cofactors (µM) by a linear fit equation. The concentration of NAD^+^, NADH, NADP^+^ and NADPH was expressed in µmol/g dry weight (DW) after converting the OD_600_ measurement to cell dry weight using the conversion factor, 1 OD_600_ = 10^9^ cells, and 1 cell dry weight = 3 x 10^-13^ g. The mean and standard deviation from three independent experiments are reported.

#### Zwf and Gnd enzyme activity assay in cell free extract

Wild type (MG1655) culture was grown overnight in LB medium and diluted 100-fold in the same medium. Cells were collected at mid-log phase (A_600_ of 0.4 to 0.6) and Glucose-6-P dehyrogenase (Zwf) and 6-phospho-gluconate dehydrogenase (Gnd) activity were assayed using cell-free extracts as described by Fraenkel and Levisohn (1967) with slight modification. Briefly, cells from 30 ml culture was collected by centrifugation at 8,000 rpm at 4°C, washed once with prechilled 0.9% NaCl and re-suspended at one-fifth of original volume in a buffer containing 0.01M Tris chloride, 0.01M MgCl_2_ and 0.001M dithiothreitol (pH-7.8). The cell suspensions were sonicated on ice for 5 minutes (30 sec on and 40 sec off) and centrifuged at 17,000g for 30 min at 4°C and the supernatants were stored at 4 °C and used on the same day of preparation. The enzyme assay was carried out by transferring 200µl of supernatant into 800µl assay buffer, to make a final 1 ml reaction mix containing 0.05M Tris chloride, 0.01M MgCl_2_, 0.001M dithiothreitol (pH 7.8), 0.2mM NADP^+^ and either 0.6mM glucose-6-phosphate (for G6PDH) or 0.4mM gluconate-6-phophate (for 6PGDH). The change in absorbance at 340 λ was followed in a Ultrospec 2100 Pro UV-VIS spectrophotometer with the cell chamber maintained at 25°C. The absorbance was followed for 3 min by recording at 30 sec intervals. A blank reaction without NADP^+^ addition was carried out to normalize for endogenous NADP^+^ reduction in cell extracts. To study the effect of ribose-5-P the assay was repeated in the presence of increasing concentration of ribose-5-P (1mM, 5mM and 10mM) in the assay buffer. The rate of increase in A_340_ was estimated by dividing the absorbance change by time in minutes. One unit of G6PDH activity was defined as the formation of 1 μmol NADPH in one min. The extinction coefficient for NADPH (6.22 mM^−1^cm^−1^) was used to calculate the units of activity in μmol min^-1^ ml^-1^. Total protein in cell extract was measured by Bradford method (1976) using bovine serum albumin as a standard and used to calculate the specific activity (units per mg protein). No enzyme activity was detected using cell free extracts from Δ*zwf* and Δ*gnd* strains, which confirmed that only the endogenous Zwf and Gnd activity was measured in the assay.

#### Sample preparation for Metabolomics

Strains with the ΔtktA::FRT ΔtktB::FRT allele were cultured overnight in the presence of shelter plasmid pAM-tktB in LB medium with ampicillin and 1mM IPTG and sub-cultured at 1:1500 dilution into same medium or LB. Cells equivalent to 0.75 A_600_ was collected by centrifugation from cultures with A_600_ between 0.3 to 0.5 and care was taken to completely remove the growth medium. Cell pellets were immediately flash freezed using liquid nitrogen and shipped in dry ice for metabolomics profiling by GC/MS-TOF at the West Coast Metabolomics centre in University of California, Davis.

## Acknowledgement

We acknowledge NBRP, Japan, for the Keio collection of single gene knock-out strains. We thank members of the Laboratory of Bacterial Genetics for suggestions and in particular the technical officer Mr. Shaffiqu for inventory and strain maintenance. This work was primarily funded by the Centre of Excellence in Microbial Biology research grant (phase-II) of the Department of Biotechnology, Government of India.

## Conflict of Interest

The authors declare no conflict of interest.

## Notes

### Competing Interest Statement

The authors have declared no competing interest.

