## supplementary figure 1 for "Evidence for role of transketolase function in the maintenance of pyridine nucleotide levels in *Escherichia coli*"

**Fig. S1A**

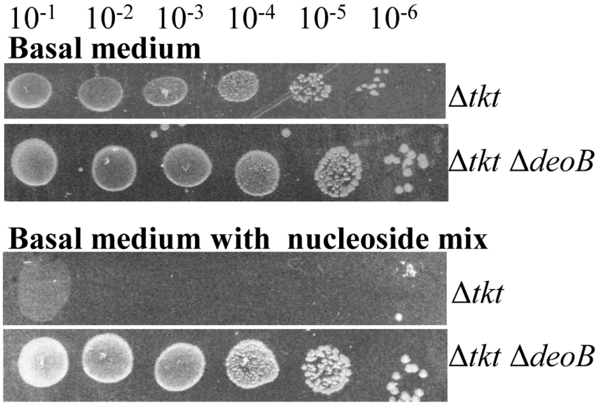


**Fig. S1A. Nucleoside catabolism by DeoB inhibits growth after transketolase depletion.** Strains with their relevant genotypes indicated were serially diluted and spotted on basal medium with or without the supplementation of nucleoside mix. Δ*tkt/*pAM-tktB (AV104); Δ*tkt* Δ*deoB/*pAM-tktB (AV155).

**Fig. S1B**

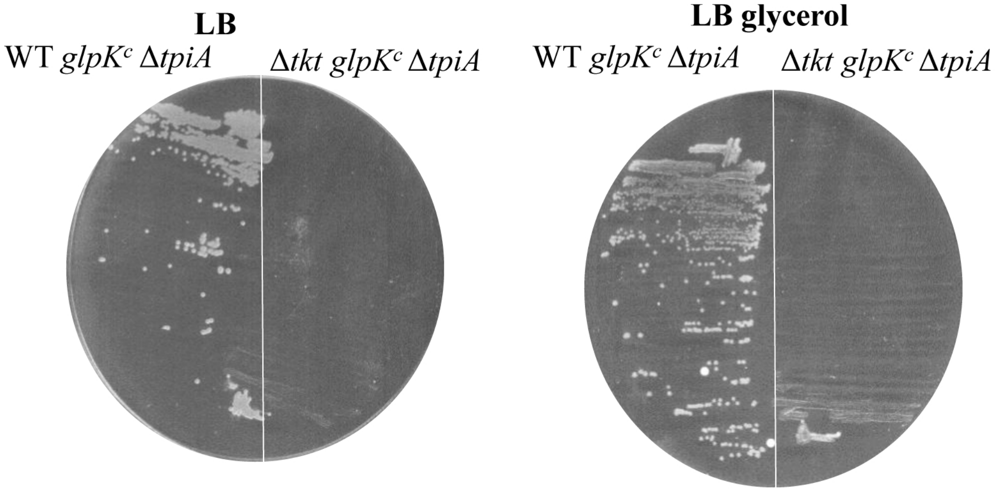


**Fig. S1B. TpiA function was required for growth rescue by glpK$^C$ allele after transketolase depletion.** Strains with their relevant genotypes indicated were streaked on LB plate with or without glycerol. Δ*tpiA glpK*$^c$ (AV38) and Δ*tkt glpK*$^c$ Δ*tpiA/*pAM-tktB (AV247).
