## supplementary figure 2 for "Evidence for role of transketolase function in the maintenance of pyridine nucleotide levels in *Escherichia coli*"

**Fig-S2**

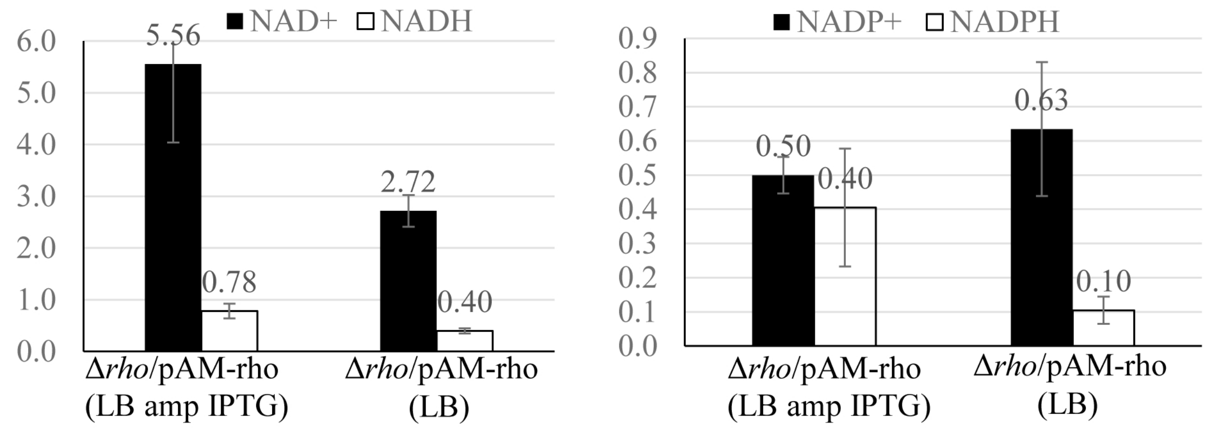


**Fig. S2. Measurement of pyridine co-factor pool before and after depletion of Rho activity.** The Δ*rho/*pAM-rho strain (GJ13490) harbouring IPTG dependent replication plasmid encoding Rho activity was grown in LB medium containing ampicillin and IPTG or in plain LB medium until depletion of Rho activity and growth arrest. Pyridine cofactor pools were measured in cultures normalized for $A_{600}$ using the cycling assay (see methods for details). The assays were carried out with three independent cultures and values reported are in µmoles/ gram dry weight. The error bars represent the standard deviation of the mean.
