## supplementary tables for "Evidence for role of transketolase function in the maintenance of pyridine nucleotide levels in *Escherichia coli*"

**Supplementary Information**

**(Tables S1 S2 & S3, Legends for Figures S1A, S1B & S2 and References)**

**Evidence for role of transketolase function in maintenance of pyridine nucleotide levels in *Escherichia coli.***

**A. Vimala^1^ and R. Harinarayanan^1^***

**^1^Laboratory of Bacterial Genetics, Center for DNA Fingerprinting and Diagnostics, Hyderabad 500039, India.**

**Keywords: Transketolase, Glycolysis, pentose phosphate pathway, Ribose-5-P, NADH, NADPH.**

**Table S1**

**List of *E.coli* K-12 strains**

| **Strain** | **Genotype** | **Source and/or reference** |
| --- | --- | --- |
| MG1655 | *E. coli* K-12 wild-type | Lab collection |
| CF14051 | MG1655 Δ*pgi*::tet | Lab collection |
| JW1841 | BW25113 Δ*zwf*::kan | (1) |
| JW2011 | BW25113 Δ*gnd*::kan | (1) |
| JW4346 | BW25113 Δ*deoB*::kan | (1) |
| JW3731 | BW25113 Δ*rbsK*::kan | (1) |
| JW 1840 | BW25113 Δ*edd*::kan | (1) |
| JW5551 | BW25113 Δ*udhA*::kan | (1) |
| JW1595 | BW25113 Δ*pntA*::kan | (1) |
| JW4347 | BW25113 Δ*deoD*::kan | (1) |
| JW3389 | BW25113 Δ*glpD*::kan | (1) |
| JW5478 | BW25113 Δ*tktA*::kan | (1) |
| JW2449 | BW25113 Δ*tktB*::kan | (1) |
| AV85 | MG1655 Δ*tktB*::FRT | (6) |
| AV88 | Δ*tktB*::FRT Δ*tktA*::kan | (6) |
| AV94 | Δ*tktB*::FRT Δ*tktA*::kan /pAM-tktB | (6) |
| AV104 | Δ*tktB*::FRT Δ*tktA*::FRT/pAM-tktB | AV94 × pCP20 |
| AV108 | Δ*tktB*::FRT Δ*tktA*::FRT Δ*pgi*::tet/pAM-tktB | AV104 × P1(CF14051) |
| AV107 | Δ*tktB*::FRT Δ*tktA*::FRT Δ*zwf*::kan/pAM-tktB | AV104 × P1(JW1841) |
| AV148 | Δ*tktB*::FRT Δ*tktA*::FRT *glpK^c^ ::*tn*10*dcm /pAM-tktB | AV104 × P1(AV105)^a^ |
| AV 150 | Δ*tktB*::FRT Δ*tktA*::FRT *glpK^c^ ::*tn*10*dcm Δ*glpD*::kan /pAM-tktB | AV148 × P1(JW3389) |
| AV 109 | Δ*tktB*::FRT Δ*tktA*::FRT Δ*edd*::kan/pAM-tktB | AV104 × P1(JW1840) |
| AV234 | Δ*tktB*::FRT Δ*tktA*::FRT *glpK^c^* ::Tn*10*dcm Δ*pgi*::tet/pAM-tktB | AV148 × P1(CF14051) |
| AV198 | MG1655 Δ*udhA*::kan | MG1655 × P1(JW5551) |
| AV309 | MG1655 Δ*udhA*::kan Δ*pgi*::tet | AV198 × P1(CF14051) |
| AV208 | Δ*tktB*::FRT Δ*tktA*::FRT Δ*udhA*::kan/pAM-tktB | AV104 × P1(JW5551) |
| AV187 | Δ*tktB*::FRT Δ*tktA*::FRT *glpK^c^ ::*tn*10*dcm Δ*udhA*::kan/pAM-tktB | AV148 × P1(JW5551) |
| AV43 | MG1655/ pACYC184 | MG1655× pACYC184 |
| AV45 | MG1655/ pACYC184-pntAB | MG1655× pACYC184-pntAB |
| AV110 | Δ*tktB*::FRT Δ*tktA*::FRT/pAM-tktB*/* pACYC184 | AV104 × pACYC184 |
| AV112 | Δ*tktB*::FRT Δ*tktA*::FRT/pAM-tktB*/* pACYC184-pntAB | AV104 × pACYC184-pntAB |
| AV130 | Δ*tktB*::FRT Δ*tktA*::FRT Δ*pntA*::kan /pAM-tktB | AV104 ×P1(JW1595) |
| AV 245 | MG1655 Δ*gapA::gapN-*cm | Centeno-Leija et al. 2013 |
| AV310 | MG1655 Δ *gapA:: gapN-*cm*/*pBR322 | AV245 × pBR322 |
| AV311 | MG1655 *Δ* *gapA:: gapN-*cm*/*pBR-udhA | AV245 × pBR322-udhA |
| AV 246 | Δ*tktB*::FRT Δ*tktA*::FRT *Δ* *gapA:: gapN-*cm /pAM-tktB | AV104 × P1(AV245) |
| AV 250 | Δ*tktB*::FRT Δ*tktA*::FRT *Δ* *gapA:: gap-*cm Δ*zwf*::kan /pAM-tktB | AV246 × P1(JW1841) |
| AV314 | Δ*tktB*::FRT Δ*tktA*::FRT *Δ* *gapA:: gapN-*cm /pAM-tktB*/* pBR322 | AV246 × pBR322 |
| AV315 | Δ*tktB*::FRT Δ*tktA*::FRT *Δ* *gapA:: gapN-*cm /pAM-tktB*/* pBR-udhA | AV246 × pBR322-udhA |
| AV155 | Δ*tktB*::FRT Δ*tktA*::FRT Δ*deoB*::kan /pAM-tktB | AV104 × P1(JW4346) |
| AV159 | Δ*tktB*::FRT Δ*tktA*::FRT Δ*deoB*::FRT /pAM-tktB | AV155 × pCP20 |
| AV134 | Δ*tktB*::FRT Δ*tktA*::FRT Δ*deoD*::FRT /pAM-tktB | AV104 × P1(JW4347) × pCP20 |
| AV160 | Δ*tktB*::FRT Δ*tktA*::FRT Δ*deoB*::FRT Δ*rbsk*::kan /pAM-tktB | AV159 × P1(JW3731) |
| AV37 | MG1655 Δ*tpiA*::kan | MG1655 × PCR (JGOtpiAFP + JGOtpiARP) |
| AV38 | MG1655 Δ*tpiA*::kan *glpK^c^::*tn*10*dcm | AV37 × P1(AV105)^a^ |
| AV247 | Δ*tktB*::FRT Δ*tktA*::FRT *glpK^c^ ::*tn*10*dcm Δ*tpiA*::kan/pAM-tktB | AV148 × P1(AV37) |
| AV186 | Δ*tktB*::FRT Δ*tktA*::FRT Δ*deoB*::FRT Δ*pntA*::kan /pAM-tktB | AV159 × P1(JW1595) |
| AV 249 | Δ*tktB*::FRT Δ*tktA*::FRT Δ*deoB*::FRT Δ*zwf*::kan /pAM-tktB | AV159 × P1(JW1841) |
| AV151 | Δ*tktB*::FRT Δ*tktA*::FRT *glpK^c^ ::*tn*10*dcm Δ*pntA*::kan /pAM-tktB | AV148 × P1(JW1595) |
| AV149 | Δ*tktB*::FRT Δ*tktA*::FRT *glpK^c^ ::*tn*10*dcm Δ*zwf*::kan /pAM-tktB | AV148 × P1(JW1841) |
| AV316 | MG1655 Δ*zwf*::kan | MG1655× P1(JW1841) |
| AV317 | MG1655 Δ*gnd*::kan | MG1655× P1(JW2011) |
| GJ13490 | MG1655 Δ*lacZYAI*::FRT Δ*rho*::kan / pAM-rho | Lab collection |

a – AV105 is a fast growing mutant obtained following λ1324 transposition in MG1655 Δ*tktB*::FRT Δ*tktA*::kan /pAM-tktB (AV94)

**Table. S2**

**List of Plasmids**

| **Plasmid** | **Description** | **Source and/or reference** |
| --- | --- | --- |
| pAM34 | IPTG dependent replication ; amp^R^ | (5) |
| pAM-tktB | pAM34 carrying *tktB* subcloned from pQE*tktB* within the EcoRI and HindIII sites | (6) |
| pCP20 | pSC101- based Ts replicon, cm^R^ amp^R^ used for invitro expression of Flp recombinase | (3) |
| pKD78 | Temperature sensitive origin , cm^R^ Designed to inactivate chromosomal gene through phage λ Red recombinase | (4) |
| pKD13 | Designed to contain an FRT-flanked kanamycin resistance (Kan) gene | (4) |
| pACYC184 | p15A origin of replication with cm^R^ and tet^R^ selectable markers | (2) |
| pACYC184-pntAB | pACYC184 carrying pntAB within EcoRI sites | This study |
| pBR322 | ColEI- based cloning vector; amp^R^ cm^R^ | (7) |
| pBR-udhA | pBR322 carrying udhA cloned within the HindIII and BamHI sites | This study |

**Table. S3**

**List of Primers**

| **Primer** | **Oligonucleotide sequence (5’-3’)** |
| --- | --- |
| JGO1324pcr | CTCGATAACTCAAAAAATACGCC |
| JGO1324seq | CAAAAGCACCGCCGGACATC |
| JGHtetA | CGCCGAAACAAGCGCTCATGAGCC |
| JGHtetB | CTATGCGCACCCGTTCTCGGAGCAC |
| JGOtpiAFP | AATCCGGCACCTGTCAGACTTAAGCCTGTTTAGCCGCTTCGTGTAGGCTGGAGCTGCTTC |
| JGOtpiARP | ATCTTCCTTTATTCGCTTATAAGCGTGGAGAATTAAAATGATTCCGGGGATCCGTCGACC |
| JGMudhAFP1 | ATAAAAGCTTTCAATTGGCTTACCCGCGATAA |
| JGMudhARP1 | ATAAGGATCCTTAAAACAGGCGGTT |
| JGMudhAFP2 | AAATTTGTTATTGCCTGCGG |
| JGMudhARP2 | CTCGGATAACCAATCACGTC |
| JGMpBR322FP | GGCGTATCACGAGGCCCTTTCG |
| JGMpBR322RP | AGGCGCCAGCAACCGCACCTGT |
| JGOspecFP | ACCCTGATAAATGCTTCAATAATATTGAAAAAGGAAGAGTATGCGCTCACGTAACTGGTCC |
| JGOspecRP | AATCAATCTAAAGTATATATATGAGTAAACTTGGTCTGACAGTTATTTGCCGACTACCTTGG |

**Legends to Supplementary Figures**

**Fig. S1A. Nucleoside catabolism by DeoB inhibits growth after transketolase depletion.** Strains with their relevant genotypes indicated were serially diluted and spotted on basal medium with or without the supplementation of nucleoside mix. Δ*tkt/*pAM*-*tktB (AV104); Δ*tkt* Δ*deoB/*pAM-tktB (AV155)

**Fig. S1B. TpiA function was required for growth rescue by *glpK*^C^ allele after transketolase depletion.** Strains with their relevant genotypes indicated were streaked on LB plate with or without glycerol. Δ*tpiA glpK^c^* (AV38) and Δ*tkt glpK^c^* Δ*tpiA/*pAM*-*tktB (AV247).

**Fig. S2. Measurement of pyridine co-factor pool before and after depletion of Rho activity.** The Δ*rho/*pAM-rho strain (GJ13490) harbouring IPTG dependent replication plasmid encoding Rho activity was grown in LB medium containing ampicillin and IPTG or in plain LB medium until depletion of Rho activity and growth arrest. Pyridine cofactor pools were measured in cultures normalized for A_600_ using the cycling assay (see methods for details). The assays were carried out with three independent cultures and values reported are in µmoles/ gram dry weight. The error bars represent the standard deviation of the mean.
